# Actin nucleotide state modulates the F-actin structural landscape evoked by bending forces

**DOI:** 10.1101/2022.06.02.494606

**Authors:** Matthew J. Reynolds, Carla Hachicho, Ayala G. Carl, Rui Gong, Gregory M. Alushin

## Abstract

ATP hydrolysis-coupled actin polymerization is a fundamental mechanism of cellular force generation. Force and actin filament (F-actin) nucleotide state in turn modulate the engagement of actin binding proteins (ABPs) to regulate actin dynamics through unknown mechanisms. Here, we show that bending forces evoke structural transitions in F-actin which are modulated by actin nucleotide state. Cryo-electron microscopy (cryo-EM) structures of ADP- and ADP-P_i_-F-actin with sufficient resolution to visualize bound solvent reveal inter-subunit interactions primarily bridged by waters which could contribute to lattice flexibility. Despite substantial ordered solvent differences in the nucleotide binding cleft, these structures feature essentially indistinguishable protein backbone conformations which are unlikely to be discriminable by ABPs. We next introduce a machine-learning enabled pipeline for reconstructing bent filaments, enabling us to visualize both continuous structural variability and side-chain level detail. Bent F-actin structures reveal major rearrangements at inter-subunit interfaces characterized by striking alterations of helical twist and deformations of individual protomers which are distinct in ADP- and ADP-P_i_-F-actin. This suggests phosphate rigidifies actin subunits to alter F-actin’s bending structural landscape. We therefore propose actin nucleotide state serves as a co-regulator of F-actin mechanical regulation, as bending forces evoke nucleotide-state dependent conformational transitions that could be readily detected by ABPs.

## Main

Forces exerted by actin polymerization power fundamental cellular processes including cell migration, organelle dynamics, and endocytosis^1^. The propulsive assembly of branched actin networks adjacent to membranes is coupled to nucleotide consumption by globular actin (G-actin) subunits, which hydrolyze bound ATP as they are incorporated into growing actin filaments (F-actin)^1^. The conversion of this chemical energy into mechanical work can be quantitatively explained by an elastic Brownian rachet mechanism^2,3^, where bending fluctuations of actin filaments contacting the membrane license subunit addition to their plus (“barbed”) ends. The elastic restoring force of these elongated filaments straightening pushes against the membrane, rectifying membrane fluctuations into directed motion. Consistent with this model, pioneering platinum replica electron microscopy studies of migrating cells’ leading edges^4^and *in vitro* assembled actin comet tails^5^, as well as more recent cryo-electron tomography studies of endocytic patches^6^and podosomes^7^, revealed substantial curvature in F-actin on the ∼200 nm length scale adjacent to surfaces experiencing polymerization forces. While actin’s nucleotide hydrolysis-coupled polymerization and exposure to bending forces are physiologically intertwined, their joint impact on F-actin structure, function, and regulation remain poorly understood.

The assembly of actin networks is coordinated by dozens of actin-binding proteins (ABPs), many of whose interactions with F-actin are modulated by actin nucleotide state and / or mechanical forces^1,8–10^ At the surface of membranes, the branched actin nucleator ARP2/3 is activated by nucleation promoting factors (NPFs), where it initiates assembly of daughter filaments. Newly incorporated actin subunits rapidly hydrolyze bound ATP in seconds, producing a metastable ADP-P_i_-F-actin state which can persist for minutes before inorganic phosphate release, resulting in long-lived ADP-F-actin^1^. During this process, retrograde flow pushes aged filaments away from the membrane^11^, resulting in a spatial gradient of F-actin nucleotide states that is thought to serve as a biochemical marker of filament age^12^. Consistent with this model, cofilin preferentially binds and severs ADP-F-actin^13,14^, providing a mechanism for selective disassembly of aged F-actin distal from the membrane. In addition to biochemical regulation coordinated by actin nucleotide state, branched actin networks are also mechanically regulated, showing increased force production against applied resistive loads^15^, a phenomenon which has been associated with remodeling of network geometry^15,16^. Both ARP2/3’s and cofilin’s interactions with F-actin are force-sensitive^17–19^, with both proteins specifically responding to filament bending^18,20^. Several other ABPs have also recently been reported to respond to forces across individual actin filaments^21–23^, yet the molecular mechanisms enabling ABPs to detect F-actin’s nucleotide state and load-bearing status remain unclear.

Actin has been reported to feature extensive structural polymorphism which is subject to regulation by polymerization, binding partners, and nucleotide state^24–26^. F-actin formation is coupled to a substantial conformational change known as the “G- to F-transition”, characterized by subunit flattening which renders the nucleotide binding cleft’s active site competent for hydrolysis^27^. In contrast to the G- to F-transition, nucleotide-state dependent conformational transitions within F-actin have been reported to be modest, with cryo-EM structures in the 3-4 Å range suggesting either localized rearrangements in the highly flexible D-loop^28,29^ or nearly identical conformations for the protein backbone^30^. ABP binding, however, can induce substantial conformational changes in F-actin, notably cofilin engagement, which stabilizes an under-twisted form of F-actin’s helical lattice under saturation binding conditions^31,32^. The lack of nucleotide-dependent conformational changes in F-actin resembling rearrangements upon engagement by nucleotide-state sensitive ABPs like cofilin challenges a traditional allosteric model for F-actin nucleotide state discrimination by ABPs.

Actin nucleotide state could instead modulate the mechanical properties of F-actin, thereby regulating F-actin’s deformability and its capacity to undergo structural transitions associated with ABP engagement^9,14^. In support of this model, classic fluorescence microscopy-based persistence length measurements demonstrated ADP-P_i_-F-actin to be stiffer than ADP-F-actin at the micron scale^33^. While to our knowledge the effects of mechanical force on F-actin structure have not previously been visualized, F-actin dynamics have been explored through coarse-grained simulations at the atomic and mesoscopic level^34^. Early modelling abstracted to the subunit level predicted coupling between filament bending and modulation of helical lattice twist (“twist-bend coupling”)^35^, an architectural perturbation which would modify inter-subunit interfaces and thus could be detected by ABPs like cofilin. In recent finer-grained simulations abstracted to rigid bodies within subdomains, ATP hydrolysis and phosphate release were found to exhibit cooperativity through the filament lattice^36^, suggesting the capacity for coupling between the propagation of lattice deformations and filament nucleotide state. However, in the absence of direct structural visualization, whether and how mechanically-evoked conformational transitions in F-actin intersect with a filament’s nucleotide state to govern ABP engagement is unknown.

## Results

### Waters form an extensive hydrogen-bonding network in F-actin’s nucleotide cleft

To investigate whether subtle F-actin conformational changes which were not detected in previous structures could explain nucleotide-state sensing by ABPs, we first sought to determine structures of ADP- and ADP-P_i_-F-actin with improved resolution. As several protocols have been reported for preparing ADP-P_i_-F-actin^9,28,30^, we validated our approach (polymerizing actin in the presence of 15 mM KH_2_PO_4_) by monitoring cofilin severing in a total internal reflection fluorescence (TIRF) microscopy assay (Methods). Consistent with previous reports, ADP-F-actin rapidly depolymerized, while ADP-P_i_-F-actin largely remained intact over a 5-minute time course. Replacing KH_2_PO_4_ with K_2_SO_4_ resulted in intermediate severing, suggesting that dramatically reduced severing of ADP-P_i_-F-actin was due to specific engagement of phosphate rather than a nonspecific effect of high salt concentration in the buffer (Supplementary Fig. 1a,b). We therefore proceeded with this preparation, employing cryo-EM and Iterative Helical Real Space Refinement (IHRSR)^37^ as implemented in RELION^38^ to determine structures of ADP- and ADP-P_i_-F-actin at 2.43 Å and 2.51 Å resolution, respectively (Fig. 1a, Supplementary Fig. 1, Methods).

**Figure 1.**
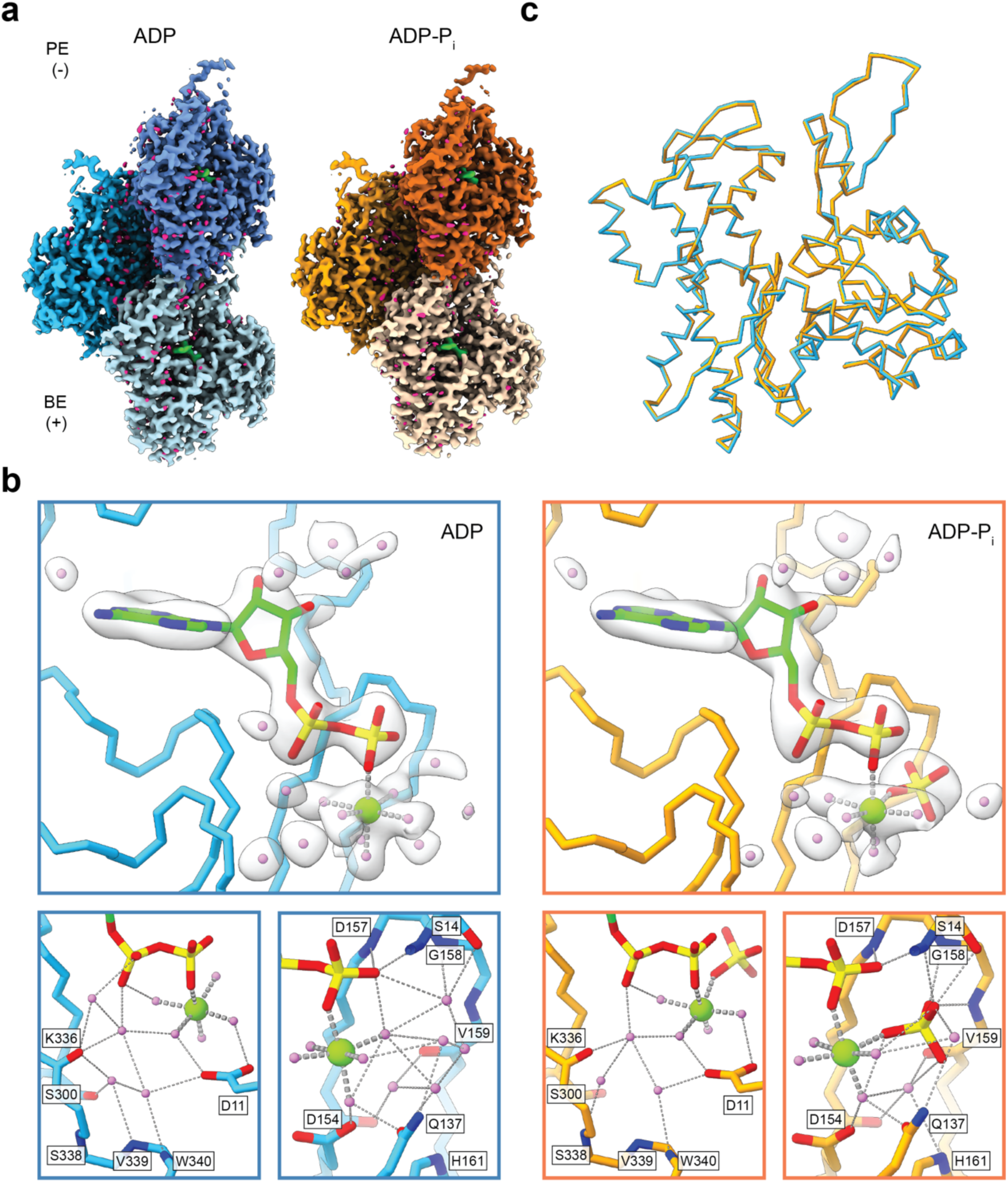
Nucleotide cleft water networks are remodeled upon phosphate release by F-actin. **(a)** Cryo-EM maps of ADP-F-actin (left, shades of blue) and ADP-P_i_-F-actin (right, shades of orange). ADP density is green, and water densities are magenta. PE: pointed end; BE: barbed end. (**b)** Top: Atomic models of the ADP- (blue) and ADP-P_i_-F-actin nucleotide clefts (orange) are shown in C_α_ representation. Carved transparent grey density is displayed for ADP (green), PO_4_^3-^ (yellow), Mg^2+^ (light green), and waters (violet). In ADP / PO_4_^3-^, nitrogen atoms are blue, oxygen atoms are red, and phosphorous atoms are yellow. Bottom panels: Backbone and side chain residues involved in putative hydrogen-bonding networks (dashed lines) are displayed and colored by heteroatom. **(c)** Superposition of individual ADP- (blue) and ADP-P_i_-F-actin (orange) protomers, displayed in C_α_ representation.

These maps provided sufficiently high-resolution views of the nucleotide cleft to accurately build and refine ADP, magnesium, inorganic phosphate, and water molecules (Fig. 1b, Supplementary Fig. 1e, Supplementary Movie 1, Methods). In both states, extensive water-mediated hydrogen-bonding networks stabilize the ligands. Strikingly, in ADP F-actin, four water molecules displace the inorganic phosphate’s oxygen tetrahedron. Minor changes in the number and arrangement of water molecules were also present in several other positions around the nucleotide cleft. The overall active site compositions and stereochemistry we observe are highly consistent with a contemporary study of F-actin nucleotide states with similar resolution by Raunser and colleagues^39^, providing confidence in interpretation of both groups’ cryo-EM maps.

Despite substantial changes in bound small molecules, differences between the conformations of actin protomers themselves, which were exquisitely resolved (Supplementary Fig. 2a), were miniscule, with an overall C_α_ RMSD of 0.202 Å (Fig. 1c). Per-residue RMSD and per-residue strain pseudo-energy analysis (a metric which highlights local deformations^40^, Methods) confirmed the general absence of notable C_α_ deviations with the exception of the D-loop in subdomain 2 (Supplementary Fig. 2c,d). This region, which is known to be flexible in F-actin^28,30,41,42^, also corresponded to the lowest resolution region of our maps, and thus differences may be partially attributable to resolution-dependent uncertainty in the atomic models. At the helical lattice level, the refined rise was slightly larger in ADP-F-actin (28.1 Å vs. 27.8 Å in ADP-P_i_-F-actin), which accumulates into detectable differences in subunit positioning at longer length scales (Supplementary Fig. 2e). Nevertheless, these structures did not reveal changes of the magnitude associated with ABP binding (e.g. cofilin^31,32^), suggesting nucleotide-state sensitivity in ABPs is unlikely to be mediated primarily by their detection of conformational changes concomitant with F-actin phosphate release.

### Waters mediate most inter-subunit contacts in F-actin

We next examined bound water molecules outside of the nucleotide cleft, reasoning solvent at inter-subunit interfaces could potentially facilitate mechanical remodeling of the filament. As is typical for helical reconstructions, the highest resolution regions of our maps were the filament cores (Supplementary Fig. 1d), which revealed a continuous solvated core aligned with the filament axis. Water positioning in this core was largely similar between ADP- and ADP-P_i_-F-actin (Fig. 2a, Supplementary Fig. 2d). Analysis of solvent-accessible pockets^43^ within the F-actin structures (Methods) indicates the filament core is accessible to the bulk solvent between protofilaments and it includes small channels to the protomers’ nucleotide clefts. This positions it to contribute to solvent-mediated lateral interactions between strands, which occur at a vertex between three subunits composed of the D-loop of subunit i, H-plug of subunit i+1, and a surface on subdomain 3 of subunit i+2 which we refer to as “site 1”. We observe only a single direct side-chain mediated interaction across strands through H-plug residue E270; however, 6 bridging waters, many of which are coordinated by backbone carbonyls, span the interface (Fig. 2b, top). While there is minor repositioning of waters between the ADP and ADP-P_i_ states, their number and coordination geometry is preserved.

**Figure 2.**
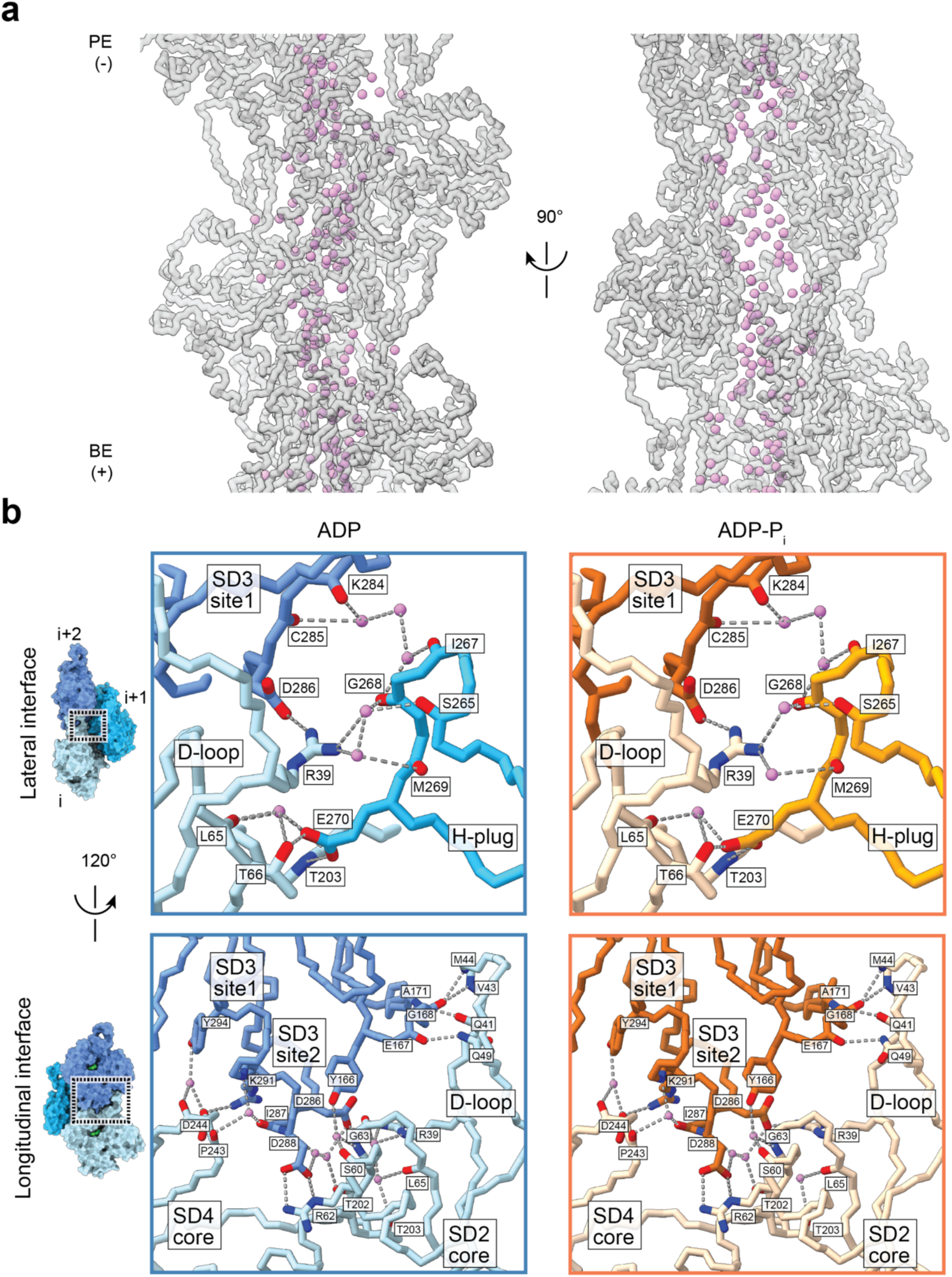
Waters mediate key longitudinal and lateral contacts in F-actin. **(a)** Water molecules (violet) contained within ADP-F-actin’s filament core. Actin subunits are shown in transparent gray mainchain representation. PE: pointed end; BE: barbed end. **(b)** Solvent-mediated contacts at longitudinal (top) and lateral (bottom) interfaces in ADP- (shades of blue) and ADP-P_i_-F-actin (shades of orange). Key sidechains and backbone atoms are displayed and colored by heteroatom. Water molecules are shown in violet, and putative hydrogen bonds are indicated with dashed lines.

We also observed ordered water molecules on the outer surface of F-actin, which contribute to extensive longitudinal interactions between protomers along the same strand (Fig. 2b, bottom). A bi-partite major interface is formed between subunit i’s subdomain 2 and a surface of subunit i+2’s subdomain 3 which we refer to as “site 2”. The D-loop forms several contacts, none of which we found to be solvent-mediated (although this may also be a limitation of local map resolution). We also observe an extensively hydrated interface on the surface of subdomain 2’s core, containing four bridging waters in the ADP-P_i_ state and one additional water in the ADP state, that features only a single direct side-chain mediated salt bridge. A minor interface is also formed between subunit i’s subdomain 4 “flap” and subunit i+2’s subdomain 3 site 1 (the same site which mediates lateral interactions), once again featuring a single side-chain mediated salt bridge, as well as two bridging waters which are equivalently positioned in the ADP and ADP-P_i_ states. In summary, we find that both lateral and longitudinal interfaces are extensively solvated, which we hypothesize could lubricate mechanical rearrangements within the filament.

### Direct visualization of bending deformations in F-actin

Actin filaments are exposed to thermal fluctuations and fluid forces^8,44^ during cryo-EM grid preparation, producing a subset of visibly bent segments (Fig. 3a) that are generally excluded during helical processing. Although other protein filament systems featuring stable curvature have recently been structurally characterized^45,46^, to our knowledge non-equilibrium bending has yet to be visualized in atomistic detail. As F-actin bending is energetically unfavorable, only a minor population of filament segments is expected to feature appreciable bending deformations. Moreover, bending is a continuum rather than a discrete conformational transition, rendering it refractory to characterization by traditional classification approaches.

**Figure 3.**
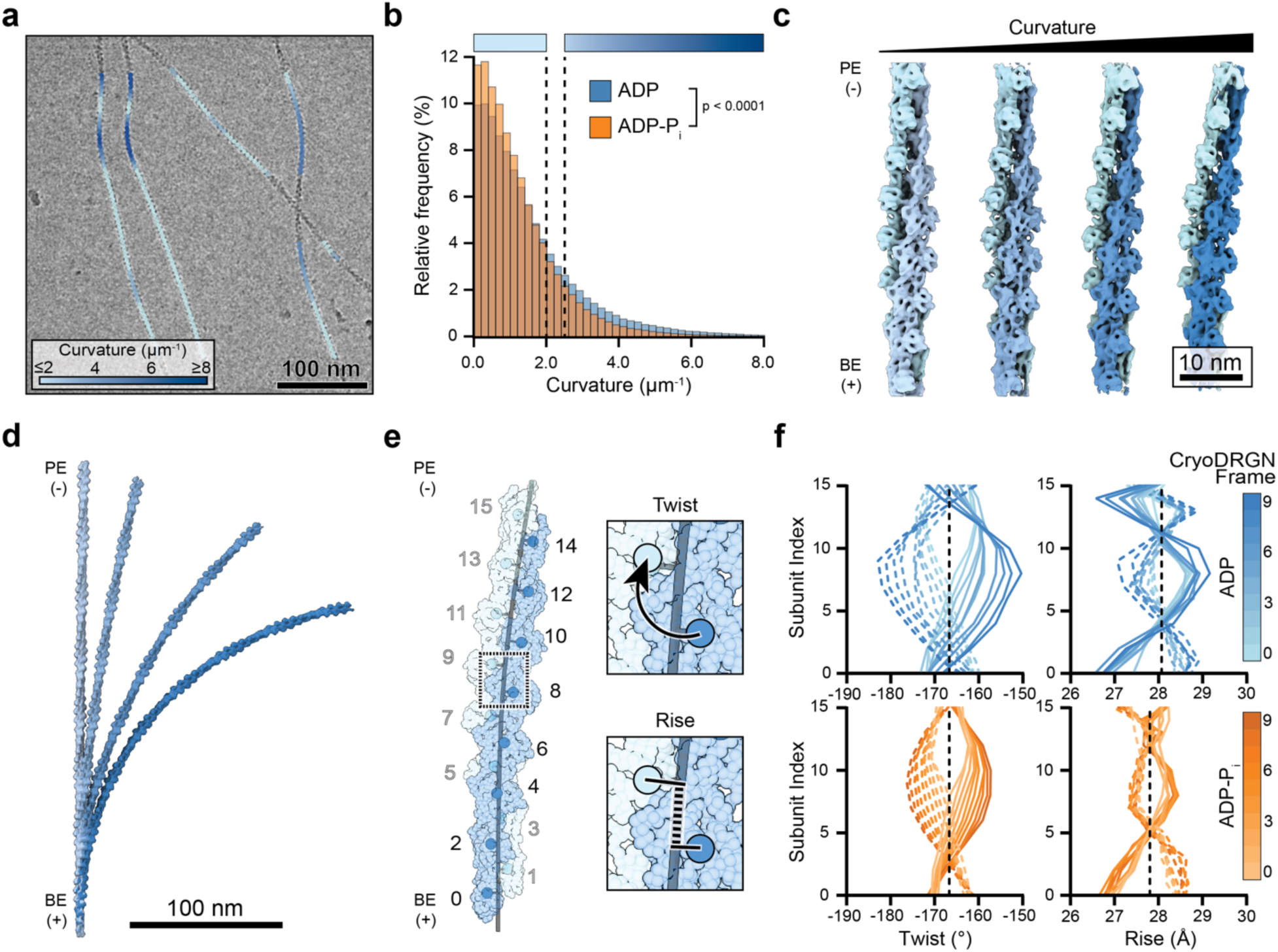
Cryo-EM reconstructions of mechanically deformed F-actin reveal bend-twist coupling. **(a)** Representative cryo-EM micrograph of ADP-F-actin filaments featuring high-curvature regions, low-pass filtered to 30 Å. Picked segments are colored by estimated curvature as indicated. **(b)** Normalized curvature histograms of ADP- (blue, n = 374,942) and ADP-P_i_-F-actin (orange, n = 470,625) filament segments, compared by Mann-Whitney test. Dashed lines indicate curvature thresholds for “straight” (≤2.0 µm^-1^) and “bent” (≥2.5 µm^-1^) segments. Color bars correspond to curvature key in a. **(c)** Helically symmetric ADP-F-actin (left map) and cryoDRGN reconstructions sampling continuous bending of ADP-F-actin (right three maps), low-pass filtered to 8 Å. Strands are colored in shades of blue. PE: pointed end; BE: barbed end. **(d)** Stitched volumes of straight and bent maps from c, aligned on the bottom 16 protomers. **(e)** Schematic of twist and rise measurements along a bent filament axis, with protomer numbering indicated. **(f)** Twist and rise measurements of cryoDRGN reconstructions sampled along major variability component. Solid and dashed curves correspond to measurements from even-to-odd and odd-to-even protomer indices, respectively.

We therefore developed a neural-network based approach to specifically identify bent F-actin segments in cryo-EM micrographs (Supplementary Fig. 3, Methods) and estimate their instantaneous in-plane curvature. As anticipated, the majority of filament segments exhibited low curvature. However, a subset of filaments featured regions of continuous curvature gradients indicative of elastic bending (Fig. 3a). Examining the distributions of segment curvatures revealed that ADP-F-actin exhibited significantly higher mean curvature than ADP-P_i_-F-actin (ADP: 1.14 µm^-1^; ADP-P_i_: 0.96 µm^-1^), with enrichment in a long tail of highly curved segments (Fig. 3b), consistent with persistence length measurements^33^. To probe the bending conformational landscapes of ADP- and ADP-P_i_-F-actin, we examined segments composed of 16 protomers featuring estimated curvature greater than an arbitrary cutoff of 2.5 µm^-1^ (hereafter “bent F-actin”). After initial model generation enabled by ab initio reconstruction as implemented in cryoSPARC^47^, and subsequent processing with RELION^38^ (Methods), we employed the recently-developed heterogeneity analysis tool cryoDRGN^48^ to generate volumes spanning the continuous bending deformations present in each dataset (Methods, Fig. 3c,d Supplementary Fig. 4a-c). These reconstructions exhibited bending that was predominantly in-plane (Fig 3d, Supplementary Fig. 4b,d), featuring central axis curvatures that fell within the distribution of estimated segment curvatures (ADP: 2.0-5.4 µm^-1^; ADP-P_i_: 3.2-4.4 µm^-1^). The resolution of cryoDRGN reconstructions cannot assessed by comparing independent half-maps, but the maps had structural features (e.g. well-resolved α-helices) characteristic of ∼8 Å resolution.

### Bending extensively remodels F-actin’s helical lattice architecture

To examine whether F-actin bending is associated with architectural remodeling of the helical lattice, we implemented a procedure for measuring the instantaneous helical parameters (rise and twist, Fig. 3e, Supplementary Movie 2) along the bent filament axis of atomic models flexibly fit into the maps with ISOLDE (Methods). This revealed striking bend-twist coupling, consistent with theoretical predictions^35^, with alternating over- and under-twisting of the strands which increased with central axis curvature. Symmetric twist deviations between the strands remain in-phase, such that the instantaneous average twist of odd / even protomers adopts the same constant value as canonical F-actin to maintain the structural integrity of the lattice. Bending evoked twist deviations up to 15°, substantially greater in magnitude than the lattice alteration induced by cofilin binding (∼5°)^31,32^. Each strand’s bend-twist relationship could be modelled analytically as a traveling wave along its component protomers (Methods), where the twist amplitude and phase within the lattice change with increasing curvature (Fig. 3f, Supplementary Fig. 5a, Supplementary Table 3). Both ADP- and ADP-P_i_-F-actin exhibit nearly identical propagation speeds in twist / protomer-index space, but ADP-F-actin features a larger amplitude coupling factor. These mathematical modeling parameters lead to the physical interpretation that a given curvature produces larger twist deviations in ADP-F-actin than ADP-P_i_-F-actin, suggesting that the ADP F-actin lattice is inherently more deformable (Supplementary Fig. 5b, Supplementary Table 3).

We also observe modest periodic deviations in rise, with a wavelength approximately half that of the twist waves (Fig. 3f), which to our knowledge has not been described in detail by theory. This manifests as an apparent standing rather than travelling wave, whose amplitude increases with curvature. While the physical basis of rise deviations is less clear, we hypothesize they result from deformations of the actin subunits themselves to accommodate twist remodeling and increased elastic energy at the subunit interfaces. Consistently, analysis of the ISOLDE models fit into the most highly curved maps revealed nucleotide-state dependent alterations of the distances and angles between subdomains of longitudinally adjacent protomers along each strand (Supplementary Fig. 5c). The most notable systematic deviations were the subdomain 1-subdomain 1, subdomain 4-subdomain 4 (Supplementary Fig. 5c, top row), and subdomain 2-subdomain 1 distances, as well as their opposing angles (Supplementary Fig. 5c, middle row). This suggests longitudinal contacts are deformed by filament bending. Inter-strand angles also separated by reference strand, suggesting the filament core water channel and lateral contacts are additionally deformed. These observations are consistent with a lubricating role for bridging waters at inter-subunit contacts. Bent ADP-P_i_-F-actin furthermore exhibited shorter average inter-subdomain distances and smaller angles, suggesting the slightly smaller average ADP-P_i_-F-actin rise could prime the filament to differentially sample F-actin’s bending conformational space. Notably, bent F-actin in both nucleotide states exhibited no clear systematic differences in the intra-subunit subdomain 2-subdomain 1-subdomain 3-subdomain 4 dihedral angle, the major parameter describing changes in subunit flattening, suggesting F-actin bending explores a distinct structural landscape from the G- to F-transition.

To validate the cryoDRGN results, we divided the ADP data into 5 approximately equally populated bins: three containing straight segments, one containing low-curvature bent segments (curvature bin 1), and one containing high-curvature bent segments (curvature bin 2). Asymmetric single-particle reconstructions (Methods) of the straight controls revealed negligible deviations from canonical F-actin’s helical parameters, while the two curved reconstructions exhibited twist and rise patterns consistent with the cryoDRGN models (Supplementary Fig. 4e,f, Supplementary Movie 3). Collectively, these data suggest bending evokes substantial conformational transitions in F-actin which are modulated by nucleotide state.

### Filament bending shears actin protomers, localizing strain to their nucleotide clefts

We next sought to visualize the detailed protomer structural deformations accompanying filament bending. Reasoning continuous conformational flexibility was limiting the resolution of the 16-protomer reconstructions, we pursued focused refinement on the central 7 subunits. Focused refinement in RELION of the central 7 subunits (Methods) substantially improved the map resolution of both bent reconstructions (ADP-F-actin = 3.6 Å, ADP-P_i_-F-actin = 3.7 Å; Fig. 4a, Supplementary Fig. 6a-c), facilitating direct atomic model building and refinement. For comparison purposes we applied this same procedure to two straight ADP-F-actin bins featuring matched numbers of segments, yielding maps with similar resolution (both 3.7 Å), which were used to build control models (Supplementary Fig. 6).

**Figure 4.**
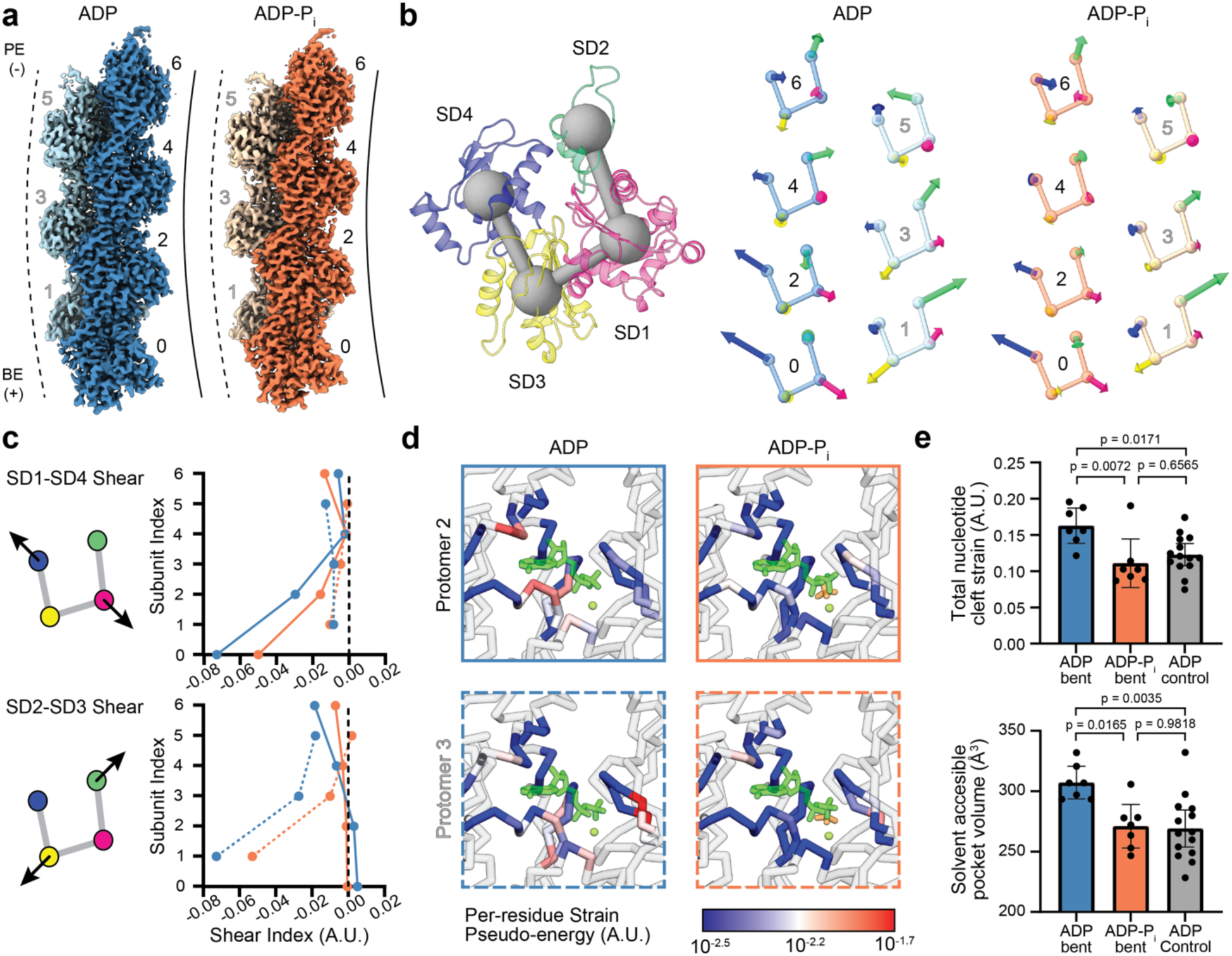
Actin nucleotide state modulates subunit shearing during filament bending. **(a)** Cryo-EM maps of bent ADP- (blue) and ADP-P_i_-F-actin (orange), colored by strand. Protomer numbering is indicated. Dashed and solid lines represent the convex and concave sides of the bent filament, respectively. PE: pointed end; BE: barbed end. **(b)** Left: ribbon representation of an individual actin protomer colored by subdomain. Spheres connected by bars indicate subdomain centroids. Right: protomer subdomain centroid diagrams with vectors (scaled 100X) indicating subdomain-averaged displacements from the corresponding helically-symmetric model (right). **(c)** Plots of subdomain shear indices (representing coordinated rearrangements) of subdomains 1 and 4 (top) and subdomains 2 and 3 (bottom). ADP: blue; ADP-P_i_: orange. Solid and dashed lines represent even (concave side) and odd (convex side) protomers, respectively. **(d)** C_α_ representation of indicated protomers’ nucleotide clefts from bent ADP- and ADP-P_i_-F-actin, colored by per-residue strain pseudo-energy. ADP (dark green), magnesium (light green), and phosphate (orange) are shown in stick representation. **(e)** Top: quantification of each protomer’s nucleotide cleft strain pseudo-energy, compared between nucleotide states and bending conditions. Bottom: equivalent quantification of solvent-accessible volume of nucleotide clefts. Bars represent means; error bars represent 95% CI. Bent, n = 7; straight n = 14, compared by one-way ANOVA with Tukey post hoc analysis; p-values corrected for multiple comparisons.

Aligning the asymmetric models with our helically symmetric models by superimposing their central protomers reveals systematic rearrangements only in the bent models, while the straight control models showed only small randomly distributed deviations, suggesting the bent structures capture conformational rearrangements rather than merely reflecting model building uncertainty at 3.6-3.7 Å resolution (Supplementary Fig. 6d). To probe the internal deformations of individual subunits, we superimposed each protomer with a subunit from the helically symmetric model of the same nucleotide state and examined subdomain-averaged C_α_ displacements (Methods, Fig. 4b, Supplementary Movie 4). This revealed complex patterns of coordinated subdomain displacements which were dependent on a protomer’s strand and position along the lattice, primarily characterized by subunit shearing rearrangements around the nucleotide binding cleft. Even-numbered protomers composing the longitudinally compressed strand on the inner (concave) surface of curvature featured opposing displacements of subdomains 1 and 4, while odd-numbered protomers composing the extended strand on the outer (convex) surface of curvature instead featured opposing displacements of subdomains 2 and 3. These rearrangements are captured by a “shear index” (Fig. 4c, the dot product of the indicated vectors), which reveals increased shearing towards the barbed end of our reconstructions, likely due to the local orientation of the lattice relative to the plane of curvature. While the pattern of subdomain displacements was highly similar between the ADP and ADP-P_i_ states (Fig. 4b), the magnitude of shearing was greater in ADP (Fig. 4c), consistent with the higher overall average curvature of bent ADP segments and greater deformability of ADP subunits.

To identify specific structural elements experiencing deformations, we once again performed per-residue strain pseudo-energy analysis^40^ (Methods), which is insensitive to rigid-body displacements of subdomains or the superposition reference frame. This revealed three primary sites of strain: the H-plug (Supplementary Fig. 7a, left), the D-loop (Supplementary Fig. 7a, right), and the nucleotide cleft (Fig. 4d). Strain in the H-plug and D-loop is consistent with their roles as primary sites of inter-subunit interactions. Strikingly, bent ADP-F-actin displayed significantly higher total strain in the amino acids adjacent to the nucleotide (distance cutoff of 7.5 Å) than bent ADP-P_i_-F-actin or the straight ADP-F-actin controls (Fig. 4d,e). Consistently, solvent-accessible volume measurements (Methods) also revealed expanded nucleotide clefts in the bent ADP-F-actin protomers relative to bent ADP-P_i_-F-actin and straight ADP-F-actin controls (Fig. 4e). These findings suggest that actin’s nucleotide cleft serves as a deformable locus coordinating mechanical rearrangements whose rigidity is dependent on phosphate occupancy, providing a structural mechanistic explanation for actin nucleotide state’s modulation of F-actin bending mechanics.

### Inter-subunit contact sites transduce filament bending to protomer deformations

As strain localized to structural elements mediating inter-subunit contacts (Supplementary Fig. 7a), we hypothesized that rearrangements at these interfaces could transduce architectural remodeling of the lattice into subunit deformations during filament bending. We therefore examined C_α_ displacements between bent and helically symmetric models superimposed in the reference frame of individual protomers, which revealed coupling between the reference subunit’s deformations and steric encroachment by its neighbors.

In bent ADP-F-actin, protomer 4 longitudinally compresses protomer 2 by wedging its subdomain 3 sites 1 and 2 into protomer 2’s nucleotide cleft, pressing down on protomer 2’s subdomain 2 core and separating its D-loop from subdomain 4 (Fig. 5a, top, Supplementary Movie 5). However, in bent ADP-P_i_-F-actin, protomers 4’s subdomain 3 sites 1 and 2 impinge in a less coordinated manner, diminishing the splitting rearrangement between protomer 2’s subdomains 4 and 2. We speculate that this is due to phosphate buttressing against subdomain 2’s intrusion into the nucleotide cleft. On the opposite strand, extension of the protomer 1-protomer 3 interface evokes a distinct response (Fig. 5a, bottom, Supplementary Movie 5). The different direction and smaller magnitude of protomer 3’s subdomain 3 site 1 incursion into protomer 1’s nucleotide cleft results in negligible displacement of protomer 1’s subdomain 4 core. Furthermore, the directional reversal of displacements in protomer 3’s subdomain 3 site 2 substantially alters the direction of D-loop repositioning. The presence of phosphate once again alters rearrangements in protomer 1’s subdomain 2 core, producing a similar splitting in the displacements of protomer 3 subdomain 3 sites 1 and 2 as that observed on the other strand, thereby modulating rearrangements in protomer 1.

**Figure 5.**
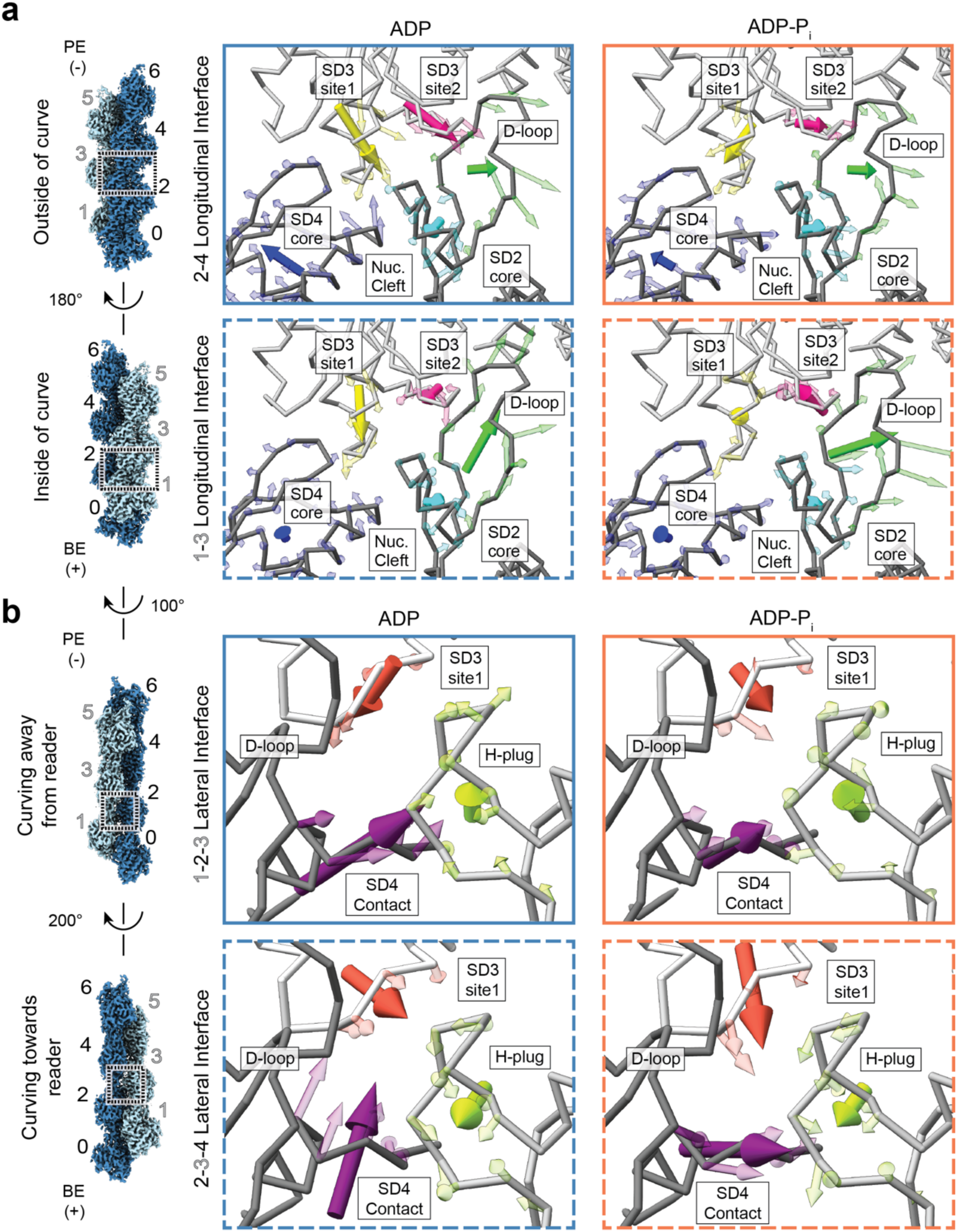
Steric encounters at inter-subunit interfaces transduce filament bending strain to protomers. **(a and b)** Longitudinal interfaces **(a)** and lateral interfaces **(b)** of bent ADP- (left) and ADP-P_i_-F-actin (right). Solid and dashed borders represent inside (concave) and outside (convex) of curve, respectively. Transparent arrows represent individual Cα displacements from helically symmetric models scaled 6X, and solid arrows show averaged displacements of indicated regions scaled 20X. PE: pointed end; BE: barbed end.

Neighboring protomers also exert steric effects at the lateral interfaces between strands (Fig. 5b, Supplementary Movie 6). In both ADP- and ADP-P_i_-F-actin, the H-plug of subunit 2 is pushed outwards normal to the direction of filament curvature, primarily due to its contact with subdomain 4 from protomer 1. In ADP-P_i_-F-actin, protomer 3’s subdomain 3 site1 also contributes to this displacement. Conversely, the H-plug of subunit 3 is pressed in towards the inside of the curve, which is driven by protomer 4’s subdomain 3 site 1 in both ADP- and ADP-P_i_-F-actin (Fig. 5b, bottom). Taken together, these data are consistent with a model in which steric incursions by neighboring protomers elicit lattice-position specific deformations in actin protomers during filament bending, with rearrangements at longitudinal interfaces in particular being modulated by the filament’s nucleotide state.

In addition to its coordination with deformations of globular subdomains, we hypothesized the well-established structural flexibility of actin’s D-loop^28,30,41,42^ could play an important role in facilitating F-actin bending. We recently reported that a short segment of the D-loop (M47-Q49) adopts a mixture of two conformations in canonical ADP-F-actin, whose prevalence is modulated by myosin motor binding^42^. Examination of the D-loop densities in our bent F-actin maps revealed considerable heterogeneity at this position, with no clear pattern based on a subunit’s lattice position or nucleotide state (Supplementary Fig. 7b). However, the density was always well-explained by the previously modelled conformations, suggesting the protomers averaged at these positions in the asymmetric reconstructions randomly sampled the plausible conformational space. This is consistent with M47-Q49 serving as flexible joint which facilitates the maintenance of inter-subunit contacts during mechanical remodeling.

## Discussion

This study reports direct structural visualization of a mechanically-regulated F-actin conformational landscape which is modulated by actin nucleotide state. The most dramatic remodeling occurs at the level of helical lattice twist, which results in substantial rearrangements at protomer-protomer contacts. We speculate F-actin bending could modulate binding interactions with numerous ABPs, as their binding sites generally span two longitudinally-adjacent protomers. Indeed, mapping the known binding sites of representative force- and nucleotide-state sensitive ABPs on bent F-actin suggests they are likely to be impacted by the structural transitions we describe here (Fig. 6a). While previous studies have postulated that phosphate release by ADP-P_i_-F-actin allosterically triggers minor rearrangements that could be detected by ABPs^28,49^, our high-resolution structures suggest this is unlikely to be the case. Our bent actin structures instead support an alternative model in which phosphate both rigidifies F-actin (consistent with previous studies)^33^ and modifies the structural landscape evoked by bending forces in a manner which could be discriminated by ABPs. Our data are also consistent with a model in which phosphate’s rigidification of canonical F-actin inhibits engagement by ABPs which must substantially deform the lattice to bind, such as cofilin^14,30,50^. The methodology we introduce here should enable future studies of ABPs in complex with mechanically deformed F-actin to dissect their joint regulation by force and actin nucleotide state.

**Figure 6.**
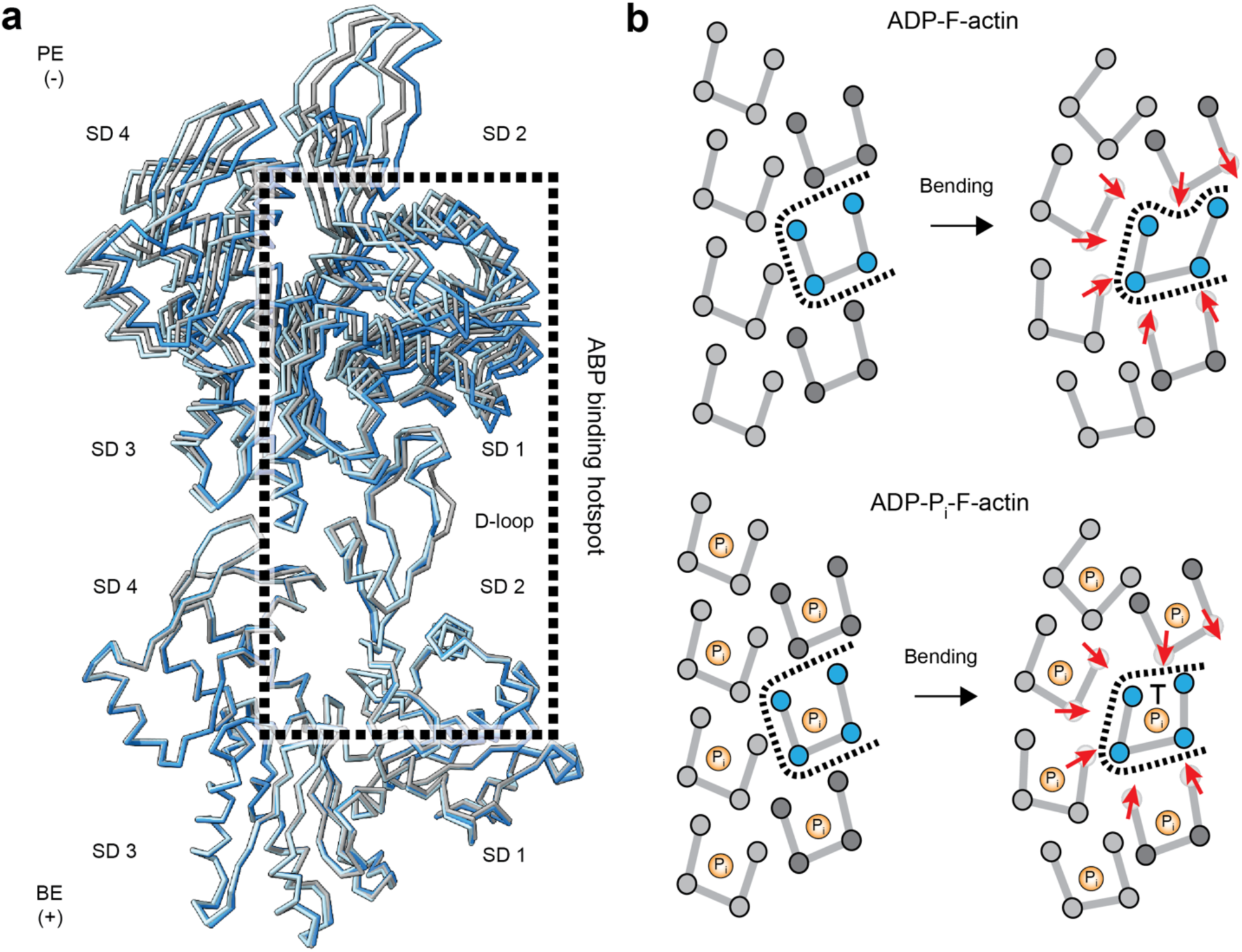
A steric boundaries mechanism for remodeling inter-subunit interfaces engaged by ABPs. **(a)** C_α_ representations of inner strand protomer 6-protomer 8 (concave, cornflower blue) and outer strand protomer 7-protomer 9 (convex, light blue) longitudinal interfaces extracted from the most curved ADP-F-actin cryoDRGN 16-protomter atomic model, superimposed on the barbed end subunit of 2 protomers extracted from the ADP-F-actin helical model (grey). Box encloses a region featuring major contacts by cofilin, coronin, ARP2/3, α-catenin, and myosin (as well as other ABPs not discussed here). Actin subdomains are indicated. BE: barbed end; PE: pointed end. (**b**) Cartoon model of steric boundaries mechanism for joint regulation of F-actin by bending forces and nucleotide state.

Our studies also provide insights into the structural mechanisms of F-actin bending, a model for investigating regulation of protein structure by mechanical force. As force is a delocalized perturbation which occurs at the filament level, it has been unclear how it is transduced into conformational remodeling of component subunits. Our data lead us to propose a “steric boundaries” conceptual model of mechanical regulation, in which repositioning of a subunit’s contacting neighbors through lattice architectural rearrangements remodels the physical space available for occupancy by the subunit, thereby inducing it to deform to minimize steric clashes. This model predicts coupling between local filament curvature, a subunit’s position in the lattice, and its resultant conformation, as we observe. The complex but stereotypical pattern of rearrangements we describe is consistent with a previous proposal that the anomalously high level of sequence conservation in actin has been evolutionarily selected to facilitate coordinated mechanical deformations of the subunit^8^. We additionally propose the extensive solvation of inter-subunit interfaces we describe here facilitates mechanical rearrangements by lubricating conformational transitions across subunit boundaries.

The steric boundaries mechanism also provides a framework for intersecting biochemical and mechanical regulation, as ligand binding interactions and chemical modifications may alter the deformation landscape of a subunit. As we show here for actin and phosphate, this does not require ligand engagement to modify the ground state conformation of the protein, a conceptual departure from traditional allosteric regulation that bears a closer resemblance to dynamic allostery^51^. However, in the steric boundaries framework, the ligand primarily operates as a co-regulator of mechanical deformations, rather than altering the intrinsic conformational fluctuations of the protein (a non-exclusive mode of regulation). Here we have shown that actin’s nucleotide binding cleft is a key site mediating subunit shearing deformations, providing an intuitive explanation for how cleft biochemical content (actin nucleotide state) modulates F-actin’s mechanics. Our observation of cleft deformations further suggests that force could in turn modulate F-actin’s biochemistry, potentially altering the kinetics of nucleotide hydrolysis and phosphate release as has previously been predicted^52^. Examination of additional F-actin nucleotide-state mimics, both experimentally and through simulations guided by the structures we report here, will be important for addressing this outstanding question. Beyond F-actin, we anticipate the steric boundaries mechanism could also facilitate joint mechanical and biochemical regulation of other multi-subunit complexes, a subject for future work.

## Acknowledgements

We gratefully acknowledge Santiago Espinosa de los Reyes (R.U.) for preparation of ADP-F-actin grids, Rama Ranganathan (U. Chicago) for strain pseudo-energy calculation pseudocode, and Hongkit Ng, Johanna Sotiris, and Mark Ebrahim from the Rockefeller University Cryo-electron Microscopy Resource Center for assistance with cryo-EM data collection. This work was funded by a National Institutes of Health grant (R01GM141044), an Irma T. Hirschl and Monique Weill-Caulier Research Award, and a Pew Biomedical Scholar Award to G.M.A. R.G. was supported by an H. Li Memorial Fellowship, and A.G.C. was supported by NIH T32 GM115327.

## Data and Materials Availability

Cryo-EM density maps and corresponding atomic models have been deposited in the PDB and EMDB with the following accession codes: Helically-symmetric ADP-F-actin (PDB: 8D13, EMDB: EMD-27114); helically-symmetric ADP-P_i_-F-actin (PDB: 8D14, EMDB: EMD-27115); asymmetric bent ADP-F-actin (PDB: 8D15, EMDB: EMD-27116); asymmetric bent ADP-P_i_-F-actin (PDB: 8D16, EMDB: EMD-27117); asymmetric straight ADP-F-actin control 1 (PDB: 8D17, EMDB: EMD-27118); asymmetric straight ADP-F-actin control 2 (PDB: 8D18, EMDB: EMD-27119). All other data needed for assessing this study’s conclusions are presented in the manuscript. Materials are available from the corresponding author (G.M.A.) without restriction.

## Code Availability

All custom code associated with this study is open-source, and it is available for download from https://github.com/alushinlab/bent_actin without restriction.

## Author Contributions

Conceptualization: MJR, GMA

Methodology: MJR, CH, AGC, RG

Investigation: MJR, CH, AGC

Formal analysis: MJR, RG, GMA

Visualization: MJR, GMA

Funding Acquisition: GMA

Supervision: GMA

Writing – Original draft: MJR, GMA

Writing – Review & editing: MJR, CH, AGC, RG, GMA

---

**Figure S1.**
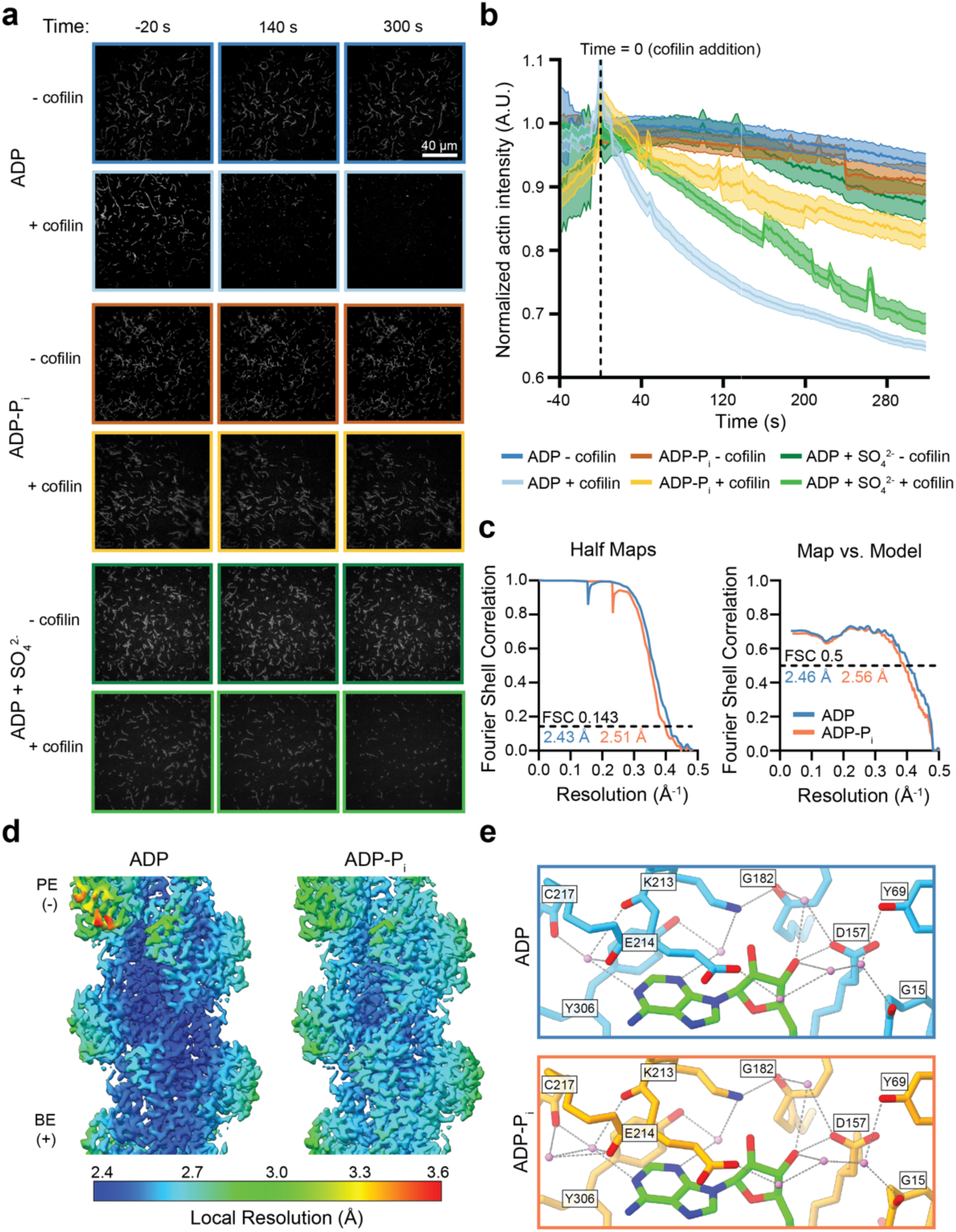
Validation of ADP-P_i_-F-actin preparation and helically symmetric reconstructions. **(a)** Representative TIRF-microscopy movie frames from cofilin severing assays. Cofilin-free controls are shown in the top row of each condition, indicated by a darker border. **(b)** Quantification of TIRF movies showing the average normalized actin channel intensity. n ≥ 3. Error margin in graph indicates +/- 95% CI. Half-lives of exponential decay at time 0 s, with 95% CI: ADP - cofilin (778 ± 24 s); ADP-P_i_ - cofilin (454 ± 14 s); ADP-sulfate - cofilin (348 ± 16 s); ADP + cofilin (50.4 ± 2.1 s); ADP-P_i_ + cofilin (177.5 ± 10.3 s); ADP-sulfate + cofilin (86.8 ± 2.5 s). **(c)** Half-map (left) and map-to-model (right) Fourier Shell Correlation (FSC) curves for helically symmetric reconstructions of ADP- and ADP-P_i_-F-actin. **(d)** Local resolution assessment of helically symmetric ADP- and ADP-P_i_-F-actin. PE: pointed end; BE: barbed end. **(e)** Potential hydrogen-bonding networks adjacent to the nucleosidyl region of ADP. Key sidechains and back bone atoms participating in hydrogen-bonding networks are displayed and colored by heteroatom.

**Figure S2.**
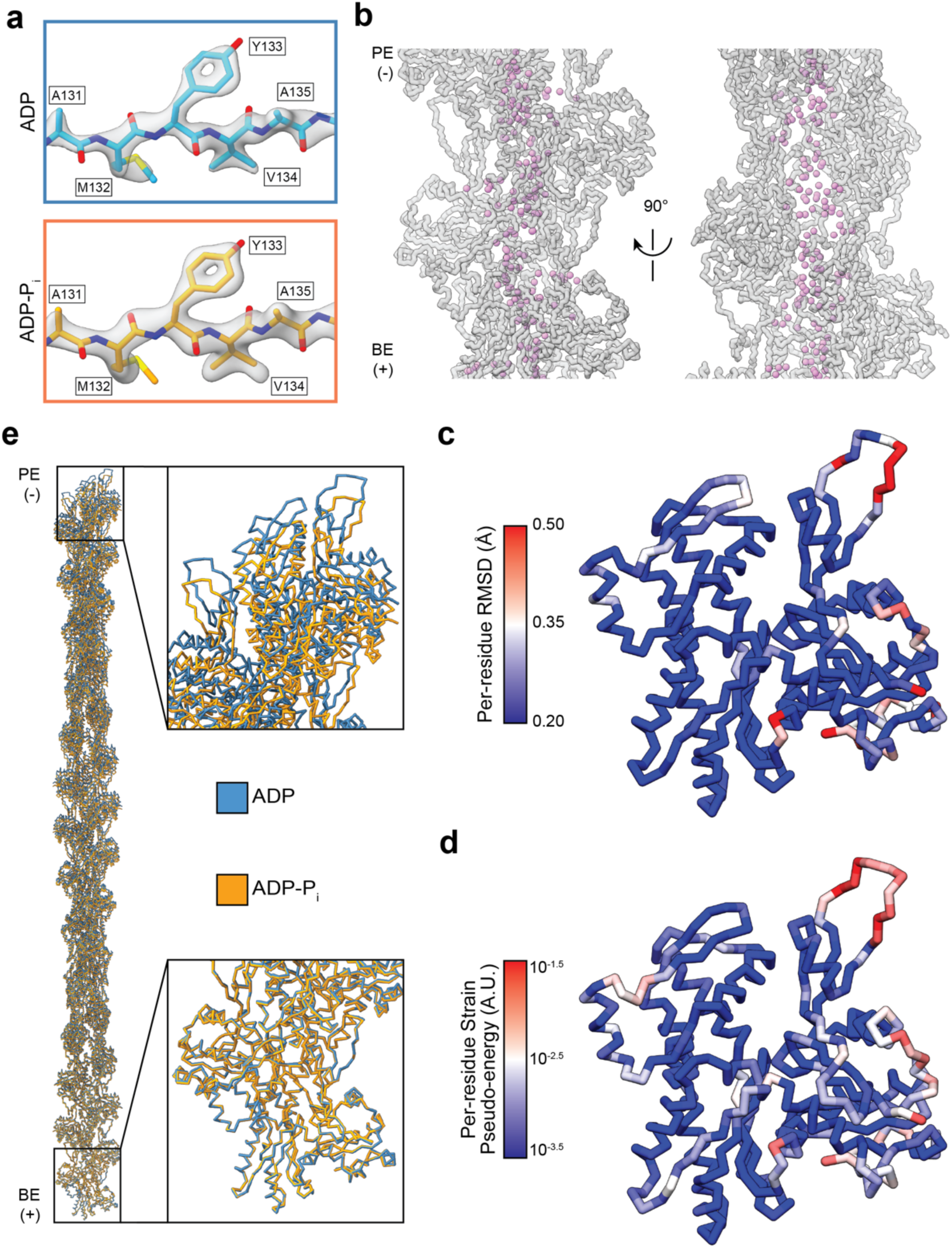
Additional analysis of helically symmetric ADP- and ADP-P_i_-F-actin models. **(a)** Example cryo-EM map density superimposed with atomic model residues A131-A135 from ADP-F-actin (top) and ADP-P_i_-F-actin (bottom). **(b)** Water molecules (violet) contained within the ADP-P_i_-F-actin filament’s core. Actin subunits are shown in transparent gray backbone representation. PE: pointed end; BE: barbed end. **(c)** Individual ADP-F-actin protomer shown in C_α_ representation, colored by per-residue RMSD between ADP- and ADP-P_i_-F-actin. **(d)** Same as c, but colored by per-residue strain pseudo-energy. **(e)** Superposition of extended 31-protomer ADP- (blue) and ADP-Pi-F-actin (orange) models, aligned at the terminal barbed end protomer.

**Figure S3.**
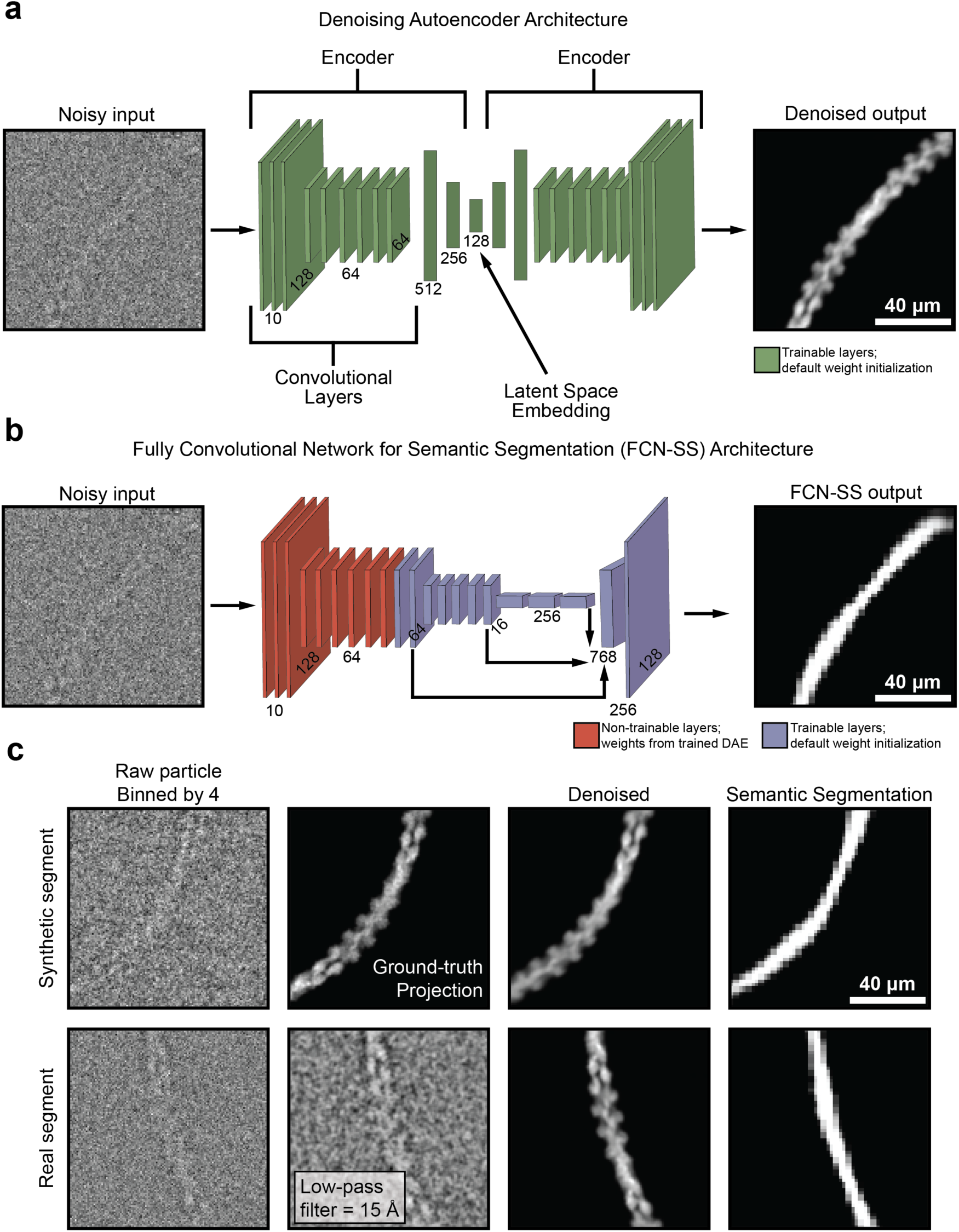
Neural network architecture and example performance. **(a)** Neural network architecture diagram for denoising auto-encoder. Example network input and output is displayed for a representative extracted segment. **(b)** Network architecture diagram for semantic segmentation fully-convolutional network. Example input and output for a representative extracted segment is shown. **(c)** Network performance on filament segments from synthetic projections (top) and experimental cryo-EM micrographs (bottom).

**Figure S4.**
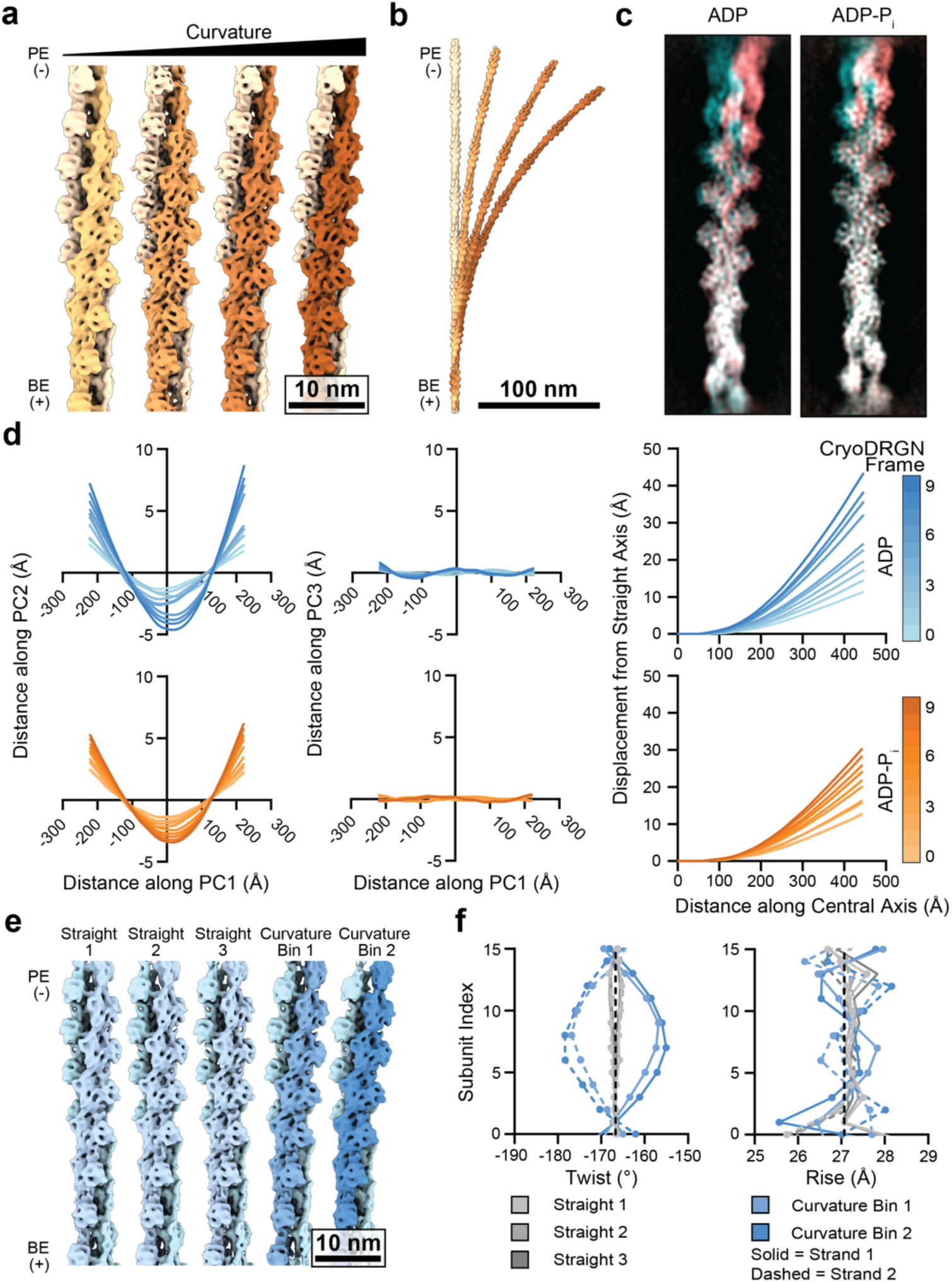
Additional analysis of filament bending deformations. **(a)** Helically symmetric ADP-P_i_-F-actin (left map) and cryoDRGN reconstructions sampling ADP-P_i_-F-actin bending (right three maps). Maps are lowpass filtered to 8 Å, and strands are colored in shades of orange. PE: pointed end; BE: barbed end. **(b)** Stitched volumes of straight and bent maps from A, aligned on the bottom 16 protomers. **(c)** Projections of zeroth (cyan) and ninth (magenta) cryoDRGN reconstructions from ADP- (left) and ADP-P_i_-F-actin (right) aligned on the bottom protomer and oriented to display maximum displacement. **(d)** Plots of central axis deviations from straight lines in ADP- and ADP-P_i_-F-actin. First and second columns show principal component analysis of the cryoDRGN reconstructions’ central axes. Third column shows displacement of the cryoDRGN reconstructions’ central axes from straight lines which were aligned to the barbed-end terminal 56 Å of the central axes. **(e)** Asymmetric reconstructions of ADP-F-actin from indicated curvature bins. **(f)** Twist and rise measurements of reconstructions from e.

**Figure S5.**
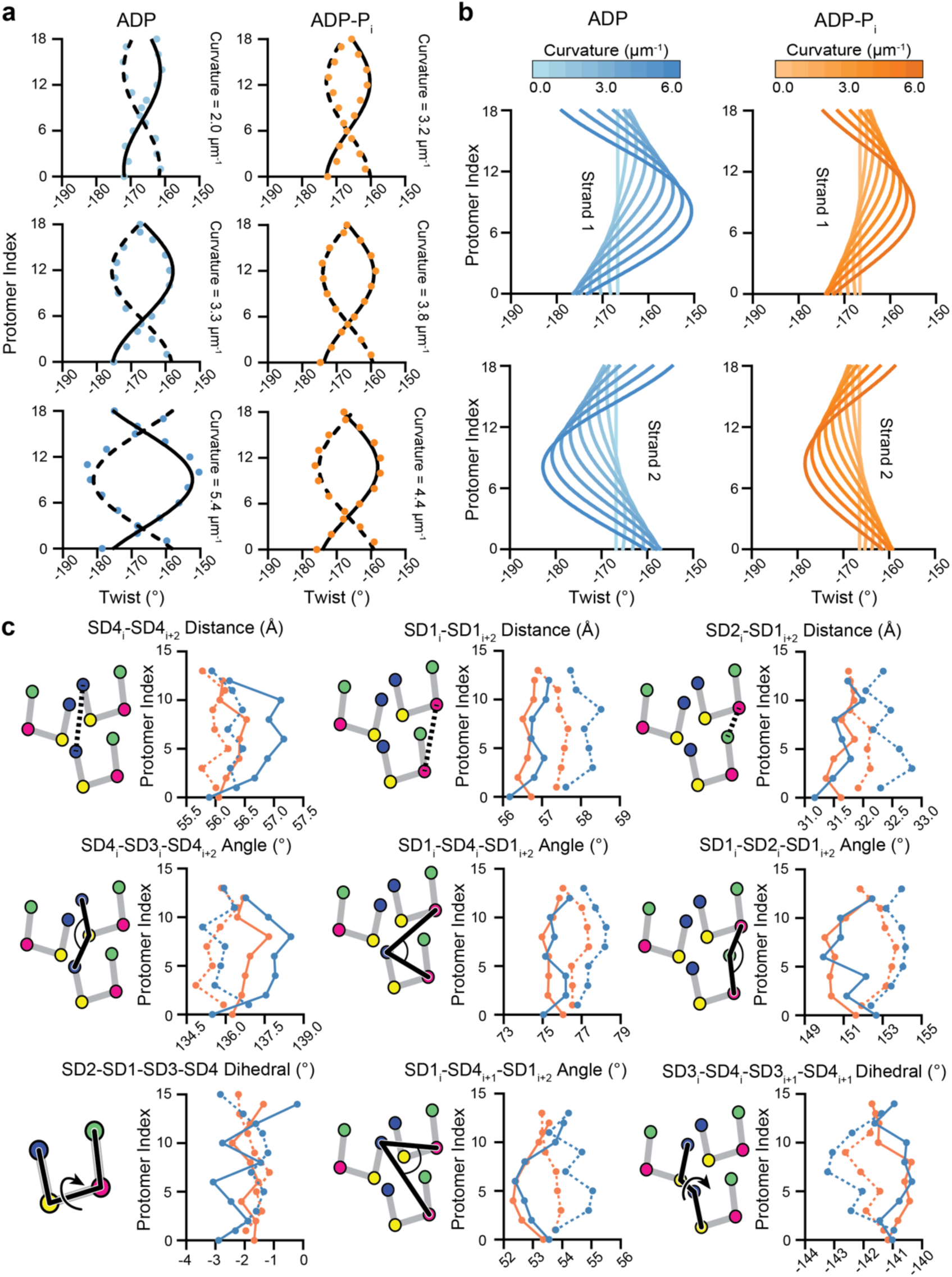
Quantitation of lattice architectural remodeling during filament bending. **(a)** Plots of traveling wave equation fits (black lines) with measured twist values (colored points) from cryoDRGN reconstructions of different curvatures. **(b)** Plots of twist traveling wave function at various curvatures, separated by strand. **(c)** Plots of intra-strand, inter-strand, and intra-protomer subdomain distances and angles from ISOLDE models of the most curved cryoDRGN reconstructions of ADP-(blue) and ADP-P_i_-F-actin (orange). Solid and dashed lines represent even (concave side) and odd (convex side) protomers, respectively.

**Figure S6.**
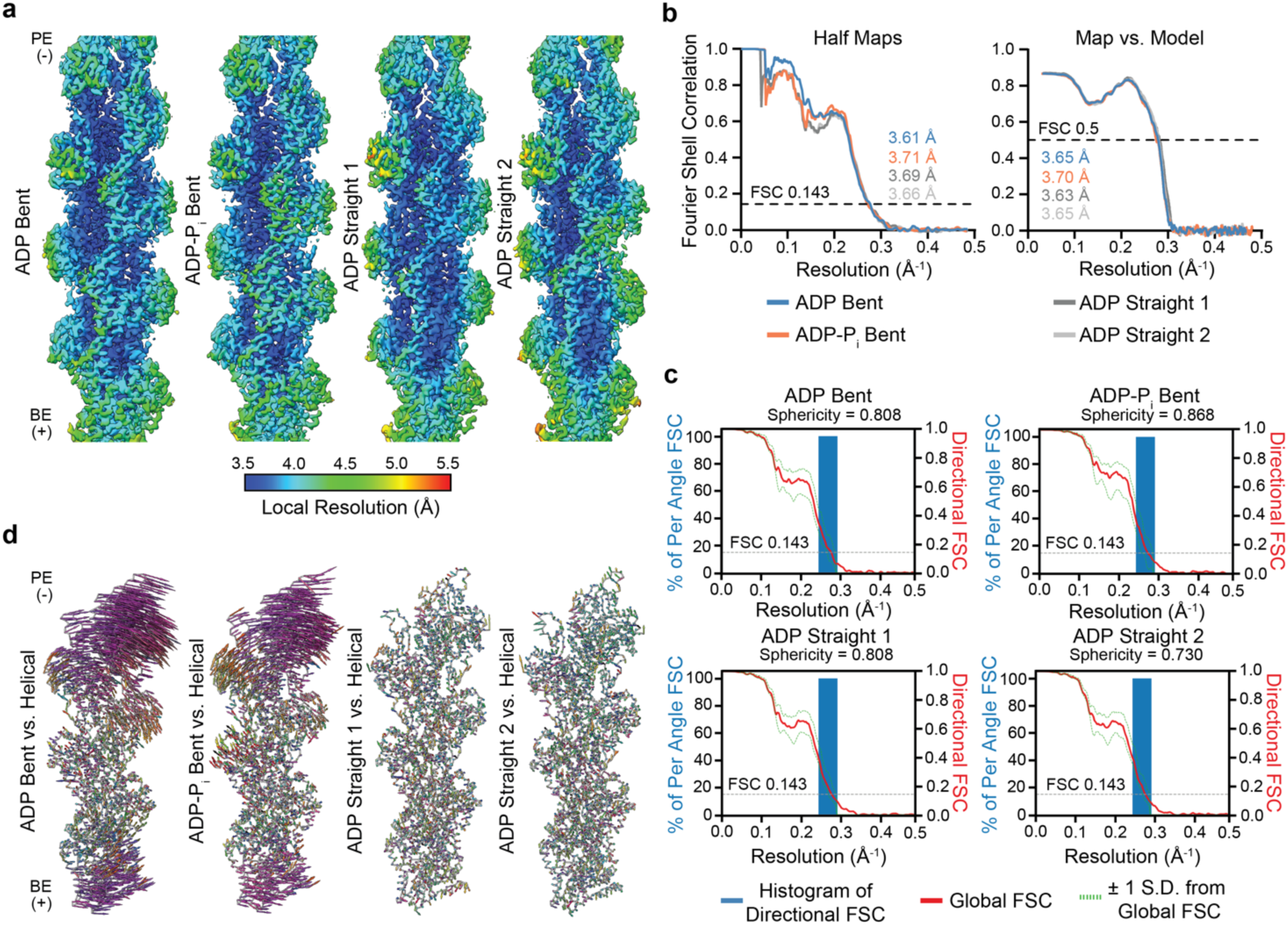
Resolution assessment and validation of bent F-actin asymmetric reconstructions. **(a)** Local resolution assessment of bent ADP-F-actin and ADP-P_i_-F-actin reconstructions, as well as two independent straight ADP-F-actin controls. PE: pointed end; BE: barbed end. **(b)** Half-map (left) and map-to-model (right) Fourier Shell Correlation (FSC) curves for asymmetric bent ADP-F-actin, ADP-P_i_-F-actin, as well as straight ADP-F-actin controls. **(c)** 3D-FSC curves for asymmetric reconstructions, which indicate equivalently isotropic resolution between bent and straight reconstructions. Dotted green lines indicate +/- 1 s.d. from average FSC. **(d)** Vector plots (colored by direction and scaled 6X) representing C_α_ displacements between helically symmetric models and those built into indicated asymmetric reconstructions, aligned on the central protomer.

**Figure S7.**
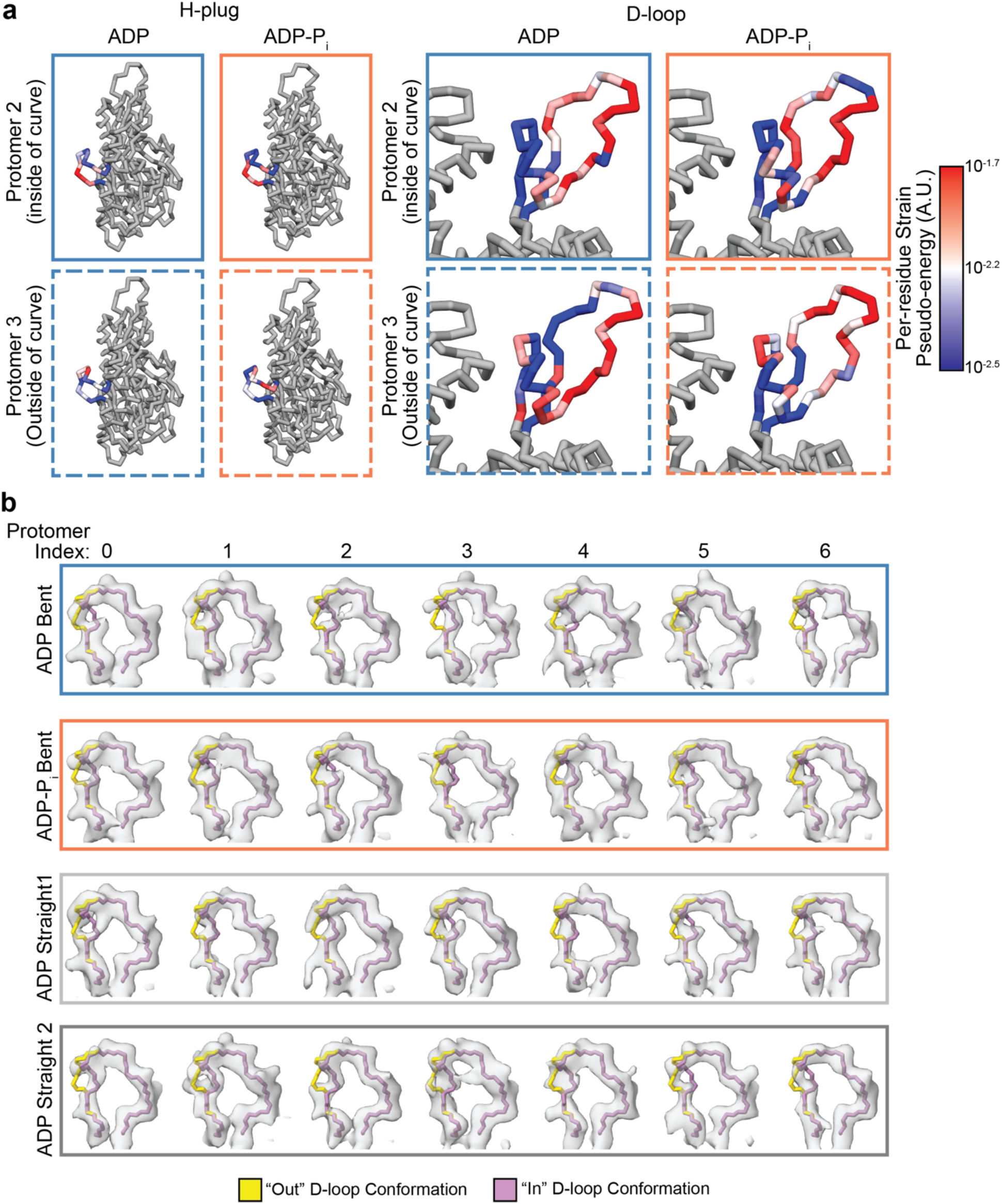
Strain and flexibility of inter-protomer contact sites in bent F-actin. **(A)** Grey C_α_ representations of indicated protomers from bent F-actin, locally colored by computed per-residue strain pseudo-energy relative to helically-symmetric models on their H-plugs (left) and D-loops (right). **(B)** D-loop heterogeneity in asymmetric F-actin maps. Local density around each of the unique D-loops from the asymmetric reconstructions are shown with a docked PDB model of ADP-F-actin (7R8V) featuring both “in” and “out” D-loop conformations.

**Supplementary Table 1.** High-resolution cryo-EM data collection, refinement, and validation statistics.

|  | ADP-F-actin<br>(EMD-27114, PDB 8D13) | ADP-Pi-F-actin<br>(EMD-27115, PDB 8D14) |
| --- | --- | --- |
| **Data collection and processing** |  |  |
| Microscope | Titan Krios | Titan Krios |
| Voltage (kV) | 300 | 300 |
| Detector | K2 Summit | K2 Summit |
| Magnification | 29,000 | 29,000 |
| Electron exposure (e<sup>-</sup>Å<sup>-2</sup>) | 60 | 60 |
| Exposure rate (e<sup>-</sup>/pixel/s) | 6 | 6 |
| Calibrated pixel size (Å) | 1.03 | 1.03 |
| Defocus range (μm) | -1.5 to -3.5 | -1.5 to -3.5 |
| Symmetry imposed | C1 | C1 |
|  | 28.07 Å rise | 27.81 Å rise |
|  | -166.69° twist | -166.64° twist |
| Initial particle images (no.) | 220,462 | 611,403 |
| Final particle images (no.) | 153,300 | 374,778 |
| Map resolution (Å) | 2.43 | 2.51 |
| FSC threshold | 0.143 | 0.143 |
| Map sharpening B factor (Å<sup>2</sup>) | -32.1 | -41.2 |
| Map resolution range (Å) | 2.40-3.42 | 2.50-3.16 |
| **Refinement** |  |  |
| Initial models (PDB ID) | 7R8V | 7R8V |
| Model resolution (Å) | 2.46 | 2.56 |
| FSC threshold | 0.5 | 0.5 |
| Model resolution range (Å) | 2.41-3.07 | 2.50-3.16 |
| Model composition | 3 actin protomers | 3 actin protomers |
| Non-hydrogen atoms | 9219 | 9249 |
| Protein residues | 1113 | 1113 |
| Ligands | 3 Mg<sup>2+</sup>, 3 ADP, 435 water | 3 Mg<sup>2+</sup>, 3 ADP, 3 PO<sub>4</sub><sup>3-</sup>, 450 water |
| **B factors (Å<sup>2</sup>)** |  |  |
| Protein | 61.15 | 66.09 |
| Ligand | 22.18 | 21.73 |
| Water | 35.34 | 48.94 |
| **R.M.S. deviations** |  |  |
| Bond lengths (Å) | 0.002 | 0.002 |
| Bond angles (°) | 0.508 | 0.506 |
| **Validation** |  |  |
| MolProbity score | 1.16 | 1.24 |
| Clashscore | 3.67 | 4.70 |
| Poor rotamers (%) | 0.64 | 0.96 |
| **Ramachandran plot** |  |  |
| Favored (%) | 99.18 | 98.91 |
| Allowed (%) | 0.82 | 1.09 |
| Disallowed (%) | 0.00 | 0.00 |
| EMRinger Score | 5.19 | 5.68 |

**Supplementary Table 2.** Asymmetric cryo-EM data collection, refinement, and validation statistics.

|  | Straight ADP-F-actin 1<br>(EMD-27118, PDB 8D17) | Straight ADP-F-actin 2<br>(EMD-27119, PDB 8D18) | Bent ADP-F-actin<br>(EMD-27116, PDB 8D15) | Bent ADP-Pi-F-actin<br>(EMD-27117, PDB 8D16) |
| --- | --- | --- | --- | --- |
| **Data collection and processing** |  |  |  |  |
| Microscope | Titan Krios | Titan Krios | Titan Krios | Titan Krios |
| Voltage (kV) | 300 | 300 | 300 | 300 |
| Detector | K2 Summit | K2 Summit | K2 Summit | K2 Summit |
| Magnification | 29,000 | 29,000 | 29,000 | 29,000 |
| Electron exposure (e<sup>-</sup>Å<sup>-2</sup>) | 60 | 60 | 60 | 60 |
| Exposure rate (e<sup>-</sup>/pixel/s) | 6 | 6 | 6 | 6 |
| Calibrated pixel size (Å) | 1.03 | 1.03 | 1.03 | 1.03 |
| Defocus range (μm) | -1.5 to -3.5 | -1.5 to -3.5 | -1.5 to -3.5 | -1.5 to -3.5 |
| Symmetry imposed | C1 | C1 | C1 | C1 |
| Initial particle images, with segment overlap (no.) | 138,415 | 138,415 | 77,559 | 124,176 |
| Final particle images, no overlap within mask (no.) | 7,833 | 7,833 | 10,753 | 14,991 |
| Map resolution (Å) | 3.69 | 3.66 | 3.61 | 3.71 |
| FSC threshold | 0.143 | 0.143 | 0.143 | 0.143 |
| Map sharpening B factor (Å<sup>2</sup>) | -47.8 | -56.0 | -64.6 | -50.0 |
| Map resolution range (Å) | 3.44-10.65 | 3.50-10.68 | 3.51-10.05 | 3.52-8.98 |
| **Refinement** |  |  |  |  |
| Initial models (PDB ID) | 8D13 | 8D13 | 8D13 | 8D14 |
| Model resolution (Å) | 3.63 | 3.65 | 3.65 | 3.70 |
| FSC threshold | 0.5 | 0.5 | 0.5 | 0.5 |
| Model resolution range (Å) | 3.44-5.87 | 3.50-5.89 | 3.51-5.53 | 3.52-5.89 |
| Model composition | 7 actin protomers | 7 actin protomers | 7 actin protomers | 7 actin protomers |
| Non-hydrogen atoms | 20,496 | 20,496 | 20,496 | 20,531 |
| Protein residues | 2597 | 2597 | 2597 | 2597 |
| Ligands | 7 Mg<sup>2+</sup>, 7 ADP | 7 Mg<sup>2+</sup>, 7 ADP | 7 Mg<sup>2+</sup>, 7 ADP | 7 Mg<sup>2+</sup>, 7 ADP, 7 PO<sub>4</sub><sup>3-</sup> |
| **B factors (Å<sup>2</sup>)** |  |  |  |  |
| Protein | 76.50 | 81.02 | 61.54 | 87.28 |
| Ligand | 43.55 | 40.50 | 28.33 | 42.44 |
| **R.M.S. deviations** |  |  |  |  |
| Bond lengths (Å) | 0.005 | 0.005 | 0.005 | 0.011 |
| Bond angles (°) | 0.664 | 0.680 | 0.678 | 0.708 |
| **Validation** |  |  |  |  |
| MolProbity score | 1.70 | 1.76 | 1.78 | 1.88 |
| Clashscore | 6.12 | 6.66 | 6.07 | 7.54 |
| Poor rotamers (%) | 0.05 | 0.00 | 0.05 | 0.00 |
| **Ramachandran plot** |  |  |  |  |
| Favored (%) | 94.70 | 94.15 | 93.10 | 92.62 |
| Allowed (%) | 5.30 | 5.85 | 6.90 | 7.38 |
| Disallowed (%) | 0.00 | 0.00 | 0.00 | 0.00 |
| EMRinger Score | 3.30 | 2.99 | 3.15 | 2.71 |

**Supplementary Table 3.**
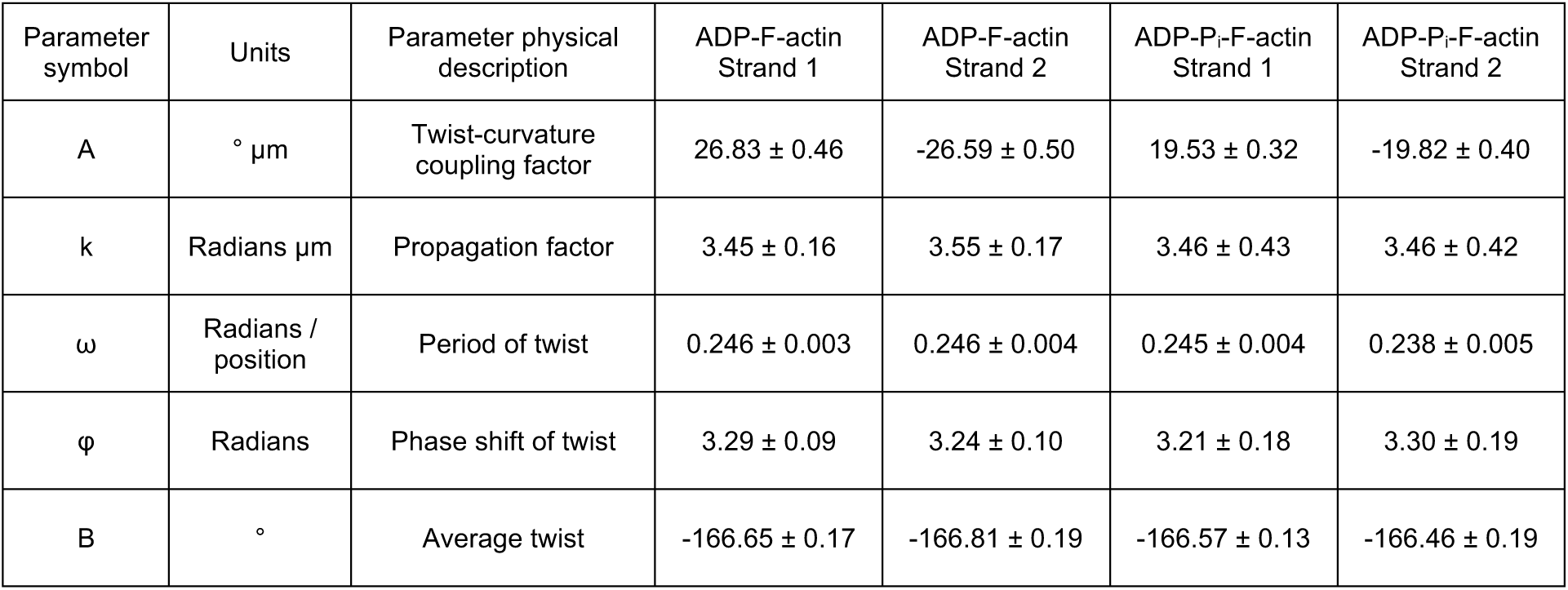
Instantaneous twist traveling wave equation parameters. Uncertainty estimates are one standard deviation error on the fit parameters.

## Supplementary Video Legends

**Supplemental Video 1. Solvated nucleotide clefts in ADP- and ADP-P_i_-F-actin**

Views of ADP- (blue) and ADP-P_i_-F-actin (orange) nucleotide clefts. Atomic models are shown in C_α_ representation. Transparent grey density from the maps is displayed. ADP: green; PO_4_^3-^: yellow; Mg^2+^ (light green), and waters (violet). In ADP / PO_4_^3-^, nitrogen atoms are blue, oxygen atoms are red, and phosphorous atoms are yellow.

**Supplemental Video 2. Morphs along bending trajectories of cryoDRGN reconstructions**

Morph of cryoDRGN reconstructions sampling continuous bending of ADP- and ADP-P_i_-F-actin low-pass filtered to 8 Å, aligned at two bottom protomers and associated helical parameters. Reconstructed ADP-F-actin strands are colored in shades of blue, and reconstructed ADP-P_i_-F-actin strands are colored in shades of orange. Rise and twist measurements are separated by strand and darken in color as curvature increases. Vertical dashed lines indicate helical parameters of canonical F-actin.

**Supplemental Video 3. Morphs of asymmetric bent and straight ADP-F-actin reconstructions**

Morph of asymmetrically reconstructed ADP maps, aligned at the central protomer. Both maps start from asymmetric, straight ADP map 1. Left morph interpolates to bent ADP map; right morph interpolates to straight ADP map 2.

**Supplemental Video 4. Morphs between straight and bent F-actin highlighting subunit shearing**

Morph of protomer subdomain shearing from helically symmetric protomer model to bent protomer models. Vectors (scaled 100X) indicate subdomain-averaged displacements.

**Supplemental Video 5. Morphs highlighting longitudinal interface deformations during bending**

Morphs of longitudinal interfaces of ADP- (left) and ADP-P_i_-F-actin (right) models from helically symmetric models to bent models. Transparent arrows represent individual C_α_ displacements from helically symmetric models scaled 6X, and solid arrows show averaged displacements of indicated regions scaled 20X.

**Supplemental Video 6. Morphs highlighting lateral interface deformations during bending**

Morphs of lateral interfaces of ADP- (left) and ADP-P_i_-F-actin (right) models from helically symmetric models to bent models. Transparent arrows represent individual C_α_ displacements from helically symmetric models scaled 6X, and solid arrows show averaged displacements of indicated regions scaled 20X.

## METHODS

### Protein preparation

Chicken skeletal muscle actin was purified as previously described^1^. Briefly, 1 g of chicken skeletal muscle acetone powder was resuspended in 20 mL of G-Ca buffer (G buffer: 2 mM Tris-Cl pH 8.0, 0.5 mM DTT, 0.2 M ATP, 0.01% NaN_3_, supplemented with 0.1 mM CaCl_2_) and mixed by inversion for 30 minutes. The suspension was centrifuged in a Beckman Ti70 rotor at 42,500 RPM (79,766 x *g*) for 30 minutes. 50 mM KCl and 2 mM MgCl_2_ was added to the supernatant, which contains G-actin monomers, to stimulate F-actin polymerization for 1 hour. 0.8 M KCl was then added, and the solution incubated for 30 minutes to facilitate dissociation of contaminants from F-actin. The solution was then centrifuged in a Ti70 rotor at 42,500 RPM (79,766 x *g*) for 3 hours. The pellet was resuspended in 2 mL of G-Ca buffer and incubated overnight. The mixture was then homogenized in a Dounce chamber for 10-15 passes, consecutively sheared through 26G and 30G needles, then dialyzed in 1 L of G-Ca buffer overnight in Spectra/Por 1 dialysis tubing (MWCO 6-8 kDa). The actin solution was then sheared through a 30G needle again before dialysis in 1 L of fresh G-Ca buffer for another day. It was then centrifuged in a Beckman Ti90 rotor at 70,000 RPM (187,354 x *g*) for 3 hours. The upper 2/3 of the supernatant was then loaded on a HiLoad 16/600 Superdex 200 column (Cytiva) for size-exclusion chromatography. Purified G-actin was maintained in G-Ca buffer at 4 °C before use.

A FLAG-GFP tagged version of human myosin VI-S1 (used to anchor actin filaments to coverslips for cofilin severing assays) was purified as described previously^2^, flash frozen in liquid nitrogen, and maintained at -80 °C. Lyophilized human cofilin 1 was purchased from Cytoskeleton (CF01) and reconstituted in MB buffer (20 mM MOPS pH 7.4, 5 mM MgCl_2_, 0.1 mM EGTA, 50 mM KCl, 1 mM DTT), then incubated overnight at 4 °C . Aliquots were then flash frozen in liquid nitrogen and maintained at -80 °C. 20 μg aliquots of lyophilized rhodamine actin (Cytoskeleton AR05) were resuspended in 18 μL of G-Ca buffer and 2 μL of milli-Q water, incubated at 4 °C for at least 1 hour, then clarified by ultracentrifugation in a Beckman TLA100 rotor at 100,000 RPM (335,400 x *g*) for 20 minutes.

### Cofilin severing assays

Glass coverslips (Corning 22 × 50 mm #1½) were cleaned for 30 minutes using 100% acetone, 10 minutes using 100% ethanol, and 2 hours using 2% Hellmanex III liquid cleaning concentrate (HellmaAnalytics) in a bath sonicator followed by rinsing with MilliQ water. The cleaned glass coverslips were coated with 1 mg/mL mPEG5K-Silane (Sigma) in a 96% ethanol, 10 mM HCl solution for at least 16 hours with rocking. After coating, the coverslips were rinsed with 96% ethanol and water, then air-dried and stored at 4°C until use.

20% rhodamine labelled ADP-F-actin was prepared by diluting unlabeled G-actin and rhodamine G-actin stocks to 0.9 µM and 0.1 µM, respectively, in KMEH buffer (50 mM KCl, 1 mM MgCl_2_, 1 mM EGTA, 10 mM HEPES pH 7.0, 1 mM DTT) supplemented with G-Mg (G Buffer + 0.1 mM MgCl_2_), for a final actin concentration of 1 µM. The mixture was incubated at 25°C for 1 hour, then 4 °C overnight. Prior to use, the F-actin was pelleted by centrifugation at 60,000 RPM (120,744 x *g*) in a TLA100 rotor for 20 minutes and resuspended in fresh KMEH in order to remove any free phosphate ions. 20% rhodamine labelled ADP-P_i_-F-actin was prepared as above, except KMEH + G-Mg was supplemented with 15 mM K_2_HPO_4_ (pH 7.0). The mixture was incubated at 25°C for 1 hour, then placed on ice and used immediately. 20% rhodamine labelled ADP-F-actin in the presence of K_2_SO_4_ was prepared identically, except 15 mM K_2_SO_4_ (pH 7.0) was substituted for K_2_HPO_4_.

For preparation of TIRF samples, a PDMS gasket (Grace Bio-Labs, Cat #103380) was placed onto the cover slip and 20 μL of 0.25 µM rigor myosin VI S1 in MB was added to the well and incubated for 2 minutes, followed by blocking with 20 μL of 0.1 % Polyvinylpyrrolidone (Sigma-Aldrich, Cat #9003-39-8) in MB for 1 minute. Next, 20 µl of 1 µM F-actin was added to the well for 30 s. The actin was then washed with 20 μL MB (for ADP-F-actin) or MB + 15 mM KH_2_PO_4_ / K_2_SO_4_ (for ADP-P_i_-F-actin / ADP-F-actin in the presence of sulfate).

TIRF movies were recorded at either a 1 s or 2 s frame rate on a Nikon H-TIRF system using a CFI Apo 60X TIRF oil immersion objective (NA 1.49), a quad filter (Chroma), and an iXon EMCCD camera (Andor). Rhodamine was excited by a 561 nm laser. Filaments were initially imaged for 2 minutes, the movie was paused, and 20 μL of 2 μM cofilin in MB (1 μM final concentration) was added to the well. The movie was then resumed and filament severing was recorded for an additional 8 minutes.

### Cofilin severing assay quantification

Movies were analyzed using custom python scripts that measured the change in filament intensity over the course of the experiments. Movie regions containing F-actin were identified and masks were generated by operating on a projection of the first 50 frames of the movie using functions in the sci-kit image python package^3^. This projection’s background (computed using a rolling ball radius of 50 pixels) was subtracted and subjected to a gaussian blur with a filter size of 2 pixels. A Li adaptive threshold was used to binarize the projection, morphological objects with an area smaller than 100 pixels were removed, and the remaining binarized image was dilated by 1 pixel. This set of masks for each movie was applied to all frames of the movie, and quantification of actin intensity was performed on a per-mask basis.

For each mask, the summed pixel intensity was measured for each frame and normalized by dividing by the 90^th^ percentile intensity. The maximum intensity was not used for normalization because the intensity often spiked with addition of buffer / cofilin at time 0 s. The intensity traces for each mask of each movie of the same experimental condition were pooled and their average was plotted (Supplemental Fig. S1).

### Cryo-EM grid preparation

ADP-P_i_-F-actin was prepared as described above (without incorporation of rhodamine actin), then diluted to 0.5 μM in KMEH + 15 KH_2_PO_4_ supplemented with 0.01% Nonidet P40 (NP40) substitute (Roche), an additive which we have found improves our ability to achieve thin vitreous ice films. 3 μl of solution was applied to a plasma-cleaned C-flat 1.2/1.3 holey carbon Au 300 mesh grid (Electron Microscopy Sciences) in a Leica EM GP plunge freezer operating at 25 °C. After 60 s incubation, the grid was blotted from the back with a Whatman no. 5 filter paper for 4 s, then flash frozen in liquid ethane.

The ADP-F-actin specimen corresponds to a pre-existing dataset described in a recent study^4^. ADP-F-actin was prepared as described above, except KH_2_PO_4_ was omitted and KMEI buffer (50 mM KCl, 1 mM MgCl_2_, 1 mM EGTA, 10 mM Imidazole pH 7.0, 1 mM DTT) + 0.01% NP40 substitute was used instead of KMEH.

### Cryo-EM data collection

ADP- and ADP-P_i_-F-actin datasets were collected on the same FEI Titan Krios operating at 300 kV equipped with a Gatan K2-Summit direct electron detector using super-resolution mode. Movies were recorded with the SerialEM software suite^5^ at a nominal magnification of 29,000X, corresponding to a calibrated pixel size of 1.03 Å at the specimen level (super-resolution pixel size of 0.515 Å / pixel). Each 10 s exposure was dose-fractionated across 40 frames, with a total electron dose of 60 e^-^ / Å^2^ (1.5 e^-^ / Å^2^ / frame), with defocus values ranging from -1.5 to -3.5 μm underfocus. For the ADP-P_i_-F-actin dataset, beam-image shift was used to collect single exposures from nine holes in a 3-by-3 grid per each stage translation. For the ADP-F-actin dataset, which has previously been reported^4^ and was reprocessed here, exposures were directly targeted using stage translations with a single exposure per hole.

## Cryo-EM Image processing

### Micrograph pre-processing

Movies were aligned with MotionCor2 using 5 × 5 patches^6^, and dose-weighting sums^7^ were generated from 2-fold binned frames with Fourier cropping, resulting in a pixel size of 1.03 Å in the images. Non-doseweighted sums were used for contrast transfer function (CTF) parameter estimation with CTFFIND4^8^.

### Synthetic dataset generation

Accurate nanoscale curvature measurements of F-actin in noisy cryo-EM micrographs requires high-quality, pixel-wise image segmentation. Traditional cross-correlation based approaches for filament particle picking use templates derived from 2D class averages or projections of a straight F-actin map. This strategy features limitations, notably that cross-correlation will be lower between straight templates and highly curved filament segments in experimental images. Moreover, discrimination of filaments from background or non-protein signal may be poor. To achieve high-quality semantic segmentation, we implemented a convolutional neural network-based approach to identify filament segments of all curvatures. While other machine-learning based pickers have recently been introduced^9–11^, to our knowledge, they do not explicitly focus on detecting or flagging instantaneous curvature within a filamentous assembly. From semantically-segmented micrographs, we identified filaments and measured their instantaneous in-plane 2D curvature.

To train the neural networks, synthetic pairs of noisy and noiseless projections that mimicked experimental data were used. Plausible three-dimensional synthetic models of F-actin bent around a circular central axis were generated using a custom python script that loaded and operated on PDB models using functions from the ProDy package^12^. Individual actin protomers were treated as rigid objects and positioned using a toroidal helix function:

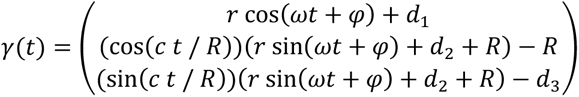

Where the parameters are defined as follows: *γ* is the position in 3D space along the toroid, *r* is the filament radius, *ω* is the average twist, *t* is the parameterized position along the helical curve, *φ* is the phase of the twist, *d_1_*, *d_2_*, and *d_3_* are the displacements of the toroid from the origin, *c* is the rise parameter, and *R* is the toroid’s radius of curvature. Note that this that this function converges to a canonical F-actin helix when the curvature is zero. Furthermore, the equation does not explicitly encode emergent architectural remodeling phenomena such as twist-bend coupling. Using this synthetic filament generation scheme, a library of 135 bent actin models of systematically varying curvature and rotation about the central filament axis were generated. These models were then converted to volume files using the PDB2MRC function in EMAN2^13^. The volumes were rotated about the phi and rot angles by a random, uniformly sampled value between 0° and 359°, and the tilt was randomly sampled from a Gaussian probability distribution centered at 90° with a standard deviation of 2.5°, then randomly translated around the box by ±250 Å and projected along the z-axis to generate a noiseless projection. A paired noisy projection was generated by adding pink noise in Fourier space, as implemented in EMAN2’s python package to generate realistic-looking synthetic data^13^. Two-channel stacks of semantic maps associated with the noisy / noiseless projection pairs were generated by low-pass filtering the noiseless projection to 40 Å and binarizing it.

### Network architecture and training

A denoising autoencoder (DAE) was trained using the architecture outlined in Supplemental Fig. 3A. Each trainable layer had a ReLU activation function, except the final layer which had a linear activation function. The negative of the cross-correlation coefficient was used as the loss function. For training, the weights were initialized using the default initialization in TensorFlow^14^. The model was trained using the Adam optimizer version of stochastic gradient descent with a learning rate of 0.00005 and minibatch size of 16 until the model converged (no improvement in validation loss for 3 epochs). Upon network convergence, the weights from the best epoch were restored. For training, 800,000 noisy / noiseless projection pairs with box sizes of 128 x 128 were generated, 90% of which were used for training and 10% for validation. Upon network convergence, the denoising autoencoder had an average cross-correlation coefficient of 0.9887 on the validation set.

After training the model as a DAE, a semantic segmentation network was trained by copying the convolutional encoding layers and weights of the DAE while adding convolutional layers. The final layer was a two-channel layer with sigmoid activation and default TensorFlow initialization. This semantic segmentation network was then trained with a learning rate of 0.001. For training, 30,000 pairs of noisy inputs and semantically segmented targets of dimension 128 x 128 and 128 x 128 x 2, respectively, were used with a minibatch size of 32; 90% of the synthetic data were used for training and 10% for validation. The loss function was binary cross-entropy, and upon network convergence the model had a loss of 0.0651 on the validation set. Example network performance on synthetic data is shown in Supplementary Fig. 3.

Models were trained on a single NVIDIA Titan XP GPU with 12 GB of VRAM. Training required approximately 1 h per epoch for the denoising autoencoder and 3 minutes per epoch for the semantic segmentation network. After this initiating this project, we have continued developing deep-learning based filament particle pickers. The architectures described here have been superseded by a U-net architecture which we have found produces better segmentation with a smaller training set in a shorter time^15^.

### Particle picking

A custom python script was used to pass images to the fully convolutional neural network for semantic segmentation (FCN-SS) and execute curvature-sensitive filament picking. Each micrograph was binned by 4 to a pixel size 4.12 Å / pixel, then 128 pixel tiles featuring 32 pixels of overlap were extracted and passed as inputs to the network. The outputs were stitched together via maximum intensity projection at the overlaps, producing a semantic segmentation map of the micrograph. These maps were then binarized using a fixed, empirically determined threshold of 0.9 and skeletonized. Branches shorter than 8 pixels were pruned, and pixels within a radius of 48 pixels from filament intersections were removed. Continuous filaments were then identified by matching tracks with common end-points, and two-dimensional splines were fit through the filaments for curvature estimation. To prevent spuriously high curvature values due to edge effects, the terminal 50 pixels of the spline were omitted from picking. From the remaining filament sections, the instantaneous curvature was measured along the spline at 56 Å intervals (corresponding to a step size the length of one protomer) and used for segment selection. For picking segments from the identified filaments for asymmetric reconstructions, a step size equivalent to two short-pitch helical rise steps (56 Å) was used for extracting segments. Filament segments with a curvature greater than or equal to 2.5 μm^-1^ were considered bent, while those with a curvature less than or equal to 2.0 μm^-1^ were considered straight. To select segments from the identified filaments for high-resolution helical reconstructions, a step size of three times the helical rise was used (83.4 Å), and segments which were members of the same filament were flagged in the output metadata (a RELION-formatted STAR file).

### Image processing of helically symmetric F-actin

We reprocessed our ADP-F-actin dataset that previously that produced a map at 2.8 Å resolution^4^, and the ADP-P_i_-F-actin dataset from this work in parallel. Our neural network-based picker eliminated intersections where filaments overlapped. After initial picking, particles were extracted without binning in 512 pixel boxes with RELION, featuring 81 Å (3 protomers) of non-overlap. Our picking scheme did not include psi (in-plane rotation) angle estimates, so an initial refinement was performed with global angular searches and a bare actin reference (EMD-24321) lowpass filtered to 35 Å. After this initial alignment, the psi angles were changed to a psi prior, all poses were removed from the metadata file, and the tilt prior was set to 90. This dataset was then subjected to a previously described F-actin cryo-EM data processing workflow^4,16^ in RELION3.1^17^, with minor modifications detailed below. Briefly, initial 2D classification was performed to remove junk particles (only 0.4% of picked particles for ADP-F-actin and 11.2% of particles for ADP-P_i_-F-actin), followed by 3D classification with alignment and five classes. For alignment, a search range of 15° about the tilt and psi priors was used, and global searches of the rot angle with an angular sampling of 7.5° was used. For ADP-F-actin, no particles were excluded at the 3D classification stage because all 5 classes were of high quality. For ADP-P_i_-F-actin, two classes comprising 23% of the remaining segments (128,533 segments total) were excluded because their helical parameters were at the border of the search range and the classes appeared abnormal. Selected particles were then subjected to unmasked 3D auto-refinement using the same angular search range described above (with local angular sampling starting at 1.7°) with helical symmetry searches as implemented in RELION3.1. This yielded a map at 4.2 Å-resolution for ADP-F-actin and 4.1 Å-resolution map for ADP-P_i_-F-actin. Post-processing was performed using a loose mask trimmed to 50% of the box size along the helical axis (“z-length”), which resulted in a resolution of 3.4 Å for ADP-F-actin and 3.5 Å for ADP-P_i_-F-actin.

Then, several iterative rounds of CTF refinement, Bayesian polishing, and 3D auto-refinement were performed. For both datasets, CTF refinement was first performed by first estimating the anisotropic magnification for each optics group. Then, defocus values were fit on a per-particle basis and astigmatism was fit on a per-micrograph basis, along with beam tilt estimation. For the ADP-P_i_-F-actin dataset, because beam-image shift was used during data collection, the data were processed in 9 optics groups. Only a single optics group was used for the ADP-F-actin dataset, which was collected with stage translations. After CTF refinement, Bayesian polishing was used to improve the data’s motion correction on a per-particle basis. The initial helical parameters for RELION’s symmetry search were updated and the mask used for post-processing was used to run another round of 3D auto-refinement. This process was repeated using a 30% z-length mask, including estimation of trefoil and 4^th^ order aberrations during CTF refinement. After the second round of particle polishing, a third round of CTF refinement was performed.

After the last iteration of CTF refinement, estimated defocus values were smoothed over the length of each continuous filament, similar to previously reported approaches^18^. Finally, a single round of masked refinement using a 30% z-length mask with local angular and translational searches was performed using solvent flattened FSCs. The final reconstructions converged with resolutions of 2.43 Å for ADP-F-actin and 2.51 Å for ADP-P_i_-F-actin.

### Image processing of bent F-actin

Selected bent segments were extracted in RELION3.0 with a box size of 512 x 512 pixels and pixel size of 1.03 Å / pixel (bin1), initially with filament overlap. To avoid reference bias, *ab initio* initial model generation was performed with cryoSPARC version 2.11.0 using the subset of ADP-F-actin segments with estimated curvature greater than 4.0 μm^-1^. Subsequent homogeneous refinement of these particles in cryoSPARC produced an asymmetric map with clear curvature. The data were then imported into RELION 3.0, and 2D classification without alignment was performed to remove junk particles, followed by unsupervised 3D classification with three classes using global angular searches. Two clearly bent, low-resolution classes curved in opposite directions and one junk class were produced. The particles in the bent classes were then subjected to supervised classification using the two bent classes as references and one straight F-actin reference as a decoy using global angular alignment. Only 0.3% of particles were assigned to the decoy, consistent with the selected segments almost exclusively featuring bent F-actin. Alignment of the two bent classes revealed them to be nearly identical but displaced by one protomer, making them appear to bend in opposite directions. Their particles were thus pooled for homogeneous refinement in cryoSPARC using global searches and the first bent class as a reference, lowpass-filtered to 30 Å. These particles were then re-imported to RELION, and underwent a 3D auto-refinement using the cryoSPARC map lowpass filtered to 10 Å as a reference, local angular searches, a loose 70% z-length mask, and solvent-flattened FSCs. This process was repeated to generate a less-bent map from segments with measured curvatures in the 2.5 μm^-1^ to 4.0 μm^-1^ range.

After demonstrating the feasibility of reconstructing bent filaments, all segments with a curvature above 2.5 μm^-1^ were then subjected to a homogeneous refinement in cryoSPARC using the highly-bent RELION refinement result as an initial reference, lowpass filtered to 30 Å. This was done for ADP-F-actin and ADP-P_i_-F-actin separately in parallel. The data were then re-imported into RELION for masked 3D auto-refinement using local searches. Successive rounds of CTF refinement, Bayesian particle polishing, and 3D auto-refinement using a 70% z-length mask were performed until resolution improvement plateaued. For the bent ADP-F-actin dataset, four rounds of CTF refinement and three rounds of Bayesian polishing were performed. For the bent ADP-P_i_-F-actin dataset, three rounds of CTF refinement and two rounds of Bayesian polishing were performed. From this stage, the data were either processed for high-resolution asymmetric analysis or continuous conformational variability analysis.

For high-resolution analysis, segment overlap within 360 pixels (corresponding to 7 protomers) was removed and the particles were subjected to a final masked (70% z-length) 3D auto-refinement with local searches and solvent-flattened FSC calculations. Resolution anisotropy of these maps was assessed with the 3DFSC server^19^. For continuous conformational variability analysis, segment overlap within the entire 512 pixel box (corresponding to 16 protomers) was removed before final masked (90% z-length) 3D auto-refinement (also with local searches and solvent-flattened FSC calculations). These segments and their assigned poses were then used for training of neural networks in cryoDRGN^20^ to assess conformational variability.

Asymmetric ADP-F-actin straight controls for the high-resolution analysis were generated using a similar method. All filament segments with a measured curvature less than or equal to 2.0 μm^-1^ were subjected to *ab initio* map generation and homogeneous refinement in cryoSPARC. They were then imported to RELION for subsequent rounds of local 3D auto-refinement, CTF refinement, and Bayesian polishing as described above. Then, all segment overlap was removed within 360 pixels, and two random subsets of sufficient size to generate maps of comparable resolution to the bent asymmetric maps were generated. These two subsets of particles underwent a final local 3D auto-refinement as described for the bent asymmetric reconstructions, and the resulting ∼3.7 Å maps were used as controls for model building and analysis.

### Variability analysis of bent F-actin

To perform variability analysis on the consensus, 16-protomer asymmetric bent reconstructions, the particles were down-sampled by 2 to a box size of 256 and a pixel size of 2.06 Å/pixel. Two cryoDRGN neural networks were trained, one for the bent ADP-F-actin dataset and one for the bent ADP-P_i_-F-actin dataset. In both cases, the network had a variational auto-encoder architecture of seven 1024-dimensional encoding and seven 1024-dimensional decoding layers with a 10-dimensional latent space. All other parameters were set to the default. The networks were trained for 40 epochs. Using four NVIDIA Titan XP GPUs the average epoch time was ∼12 minutes. Principal component analysis of the individual particle embeddings in the trained 10-dimensional latent space revealed the major variability present in the dataset to be flexible conformational heterogeneity due to bending deformations. Predicted reconstructions sampled along this trajectory in the latent space were used in subsequent analysis.

## Atomic model building and analysis

### Model building and refinement of high-resolution, helically symmetric F-actin

Prior to model building and refinement, maps were subjected to density modification with phenix.resolve_cryo_em^21^, using only the maps as inputs, then resampled on to a grid featuring 0.2575 Å voxels via four-fold Fourier un-binning with the program resample.exe (distributed with FREALIGN^22^.

Our previous model of bare ADP-F-actin (PDB: 7R8V) was copied and rigid-body fit into the central three protomers in the ADP- and ADP-P_i_-F-actin maps using UCSF Chimera^23^. Atomic models for each nucleotide state were then built and refined independently. Manual adjustments were made to the central protomer using Coot^24^, and the other two chains were replaced with this updated protomer. These models, containing three actin protomers with associated ADP, and Mg^2+^ (and PO_4_^3-^) ligands were refined using phenix real-space refinement^25^ with NCS restraints. After real-space refinement in phenix, initial solvent water molecules were placed using phenix.douse^26^ with the mean scale parameter set to 0.4. Approximately 140 waters per protomer were initially placed with this automatic function. The maps and models were then manually inspected in Coot, and waters were added or pruned. As the map resolution decreased radially from the filament core, phenix.douse was unable to reliably detect water peaks in all map regions using a single threshold. After manual adjustments, all waters outside a central slab 28 Å along the filament axis (the approximate span of a single helical rise) were deleted. The waters within the slab were then symmetrized to make an 84 Å-long slab containing waters, and two protomer chains were added to the model to fully satisfy all neighbor contacts for the central protomer. Water molecules were then associated with the closest protomer. This central protomer was copied twice and each protomer with its ligands were then fit as rigid bodies into the map to form a new trimer model. Each water molecule was then manually inspected in Coot and adjusted to fit into the map density if needed. A final phenix real-space refinement was performed with NCS restraints on the protein chains but not the solvent waters. A summary of validation statistics is provided in Supplementary Table 1.

### Model building and refinement of 16-protomer models in cryoDRGN reconstructions

From each cryoDRGN frame, 16 copies of the corresponding helically-symmetric central actin protomer model were rigid-body fit into the central 16 protomer sites in the map and combined into a single model. The map and initial model were then adjusted using the molecular dynamics flexible fitting^27^-based modelling software ISOLDE^28^ implemented in ChimeraX^29^, using secondary structure distance and torsional restraints. Due to the relatively low resolution of the cryoDRGN predictions (by visual inspection estimated to be ∼8 Å), the map weight was reduced to 10% of the automatically determined weight. The simulation temperature was set to 120 K, and the flexible fitting simulation was run for five real-time clock minutes before lowering the simulation temperature to 0 K and stopping it. These models were subsequently used for measurement of helical parameters and subdomain distance / angle measurements.

### Model building and refinement of asymmetric bent F-actin

Prior to model building, maps were subjected to standard post-processing with RELION. Models were built into the ∼3.6 Å asymmetric reconstructions by rigid-body fitting the central protomer from the corresponding helically-symmetric model into each of the seven central protomer sites in the map along the filament length. Initial, large-scale adjustments to the model were made using ISOLDE. For each condition, the map and model were loaded without applying any restraints, and the simulation temperature was set to 120 K. The simulation was then run for 5 minutes before lowering the simulation temperature to 0 K and ending the simulation. These models were then subjected to PHENIX real-space refinement without using NCS. A summary of validation statistics is provided in Supplementary Table 2.

### Multi-protomer F-actin model generation and superposition for rise analysis

The extended 31-protomer helically-symmetric actin filament models for ADP- and ADP-P_i_-F-actin were generated using UCSF Chimera as previously described^4^. Starting from the modelled actin trimer, a copy was generated and the two terminal protomers on the barbed end were superimposed on the two terminal protomers on the pointed end. These two protomers of the newly-generated trimer were deleted, and the remaining protomer was combined with the original model, extending it by one protomer. This process was repeated iteratively until a 31-protomer filament was generated.

### Water central pore and nucleotide pocket analysis

The CASTp web server^30^ was used to identify continuous solvent-accessible pockets within the high-resolution helically-symmetric F-actin structures. To eliminate boundary effects, each model was extended to five protomers as described above. Using an initial probe size of 1.4 Å revealed a solvent-accessible core that connected via narrow channels to the nucleotide pocket and broader channels to the filament’s exterior and bulk solvent. Increasing the probe size to 1.6 Å isolated the central solvent channel from these pockets. Waters contained within this discrete pocket at the filament’s core are displayed in Fig. 2 and Supplementary Fig. 2.

The CASTp server was also used with a probe size of 1.4 Å to measure the volume of the solvent-accessible nucleotide pocket in the asymmetric, 7-protomer F-actin models. The nucleotide pocket volume of individual protomers from each model was measured to separate the nucleotide pocket from the filament’s central core pocket.

### Rise / twist / curvature measurements and traveling wave analytical model

Rise and twist were measured along deformed actin filament axes using custom python scripts, implementing an approach similar to previously described helix deformations^31–33^. First, a central axis was defined using a 3D spline fit. To minimize edge effects, the model was extended by copying the model twice, aligning the terminal three subunits of one copy’s barbed end with the terminal three subunits of the original model’s pointed end and then aligning the terminal three subunit’s of the other copy’s pointed end with the terminal three subunits of the original model’s barbed end. The overlapping subunits from the copied models were deleted and the final, 42-protomer model used to define the central spline, with edge effects sufficiently minimized, was generated.

To define the 3D spline for the each filament’s central axis, an iterative, orientation-independent approach was implemented using a set of waypoints. The initial 41 waypoints were defined as the set of 3D coordinates corresponding to the centroid of 2 consecutive subunits in a rolling window along the filament. Then, a 3D cubic spline with a natural boundary condition was fit through the set of waypoints to generate the initial filament axis. A set of line segments with a length equal to the filament’s radius and one end positioned at each subunit’s centroid was aligned to minimize the free end’s distance from the spline. The waypoints were then updated to become the Euclidean average of two of these consecutive free ends. The process of updating the 3D cubic spline, defining new line segment extensions from the subunit centroids, and updating waypoints was repeated 500 times to obtain the final central axis spline.

Rise was measured by computing the distance travelled along the path of the central axis spline between protomer centroids. For each protomer, the point on the central axis that was the closest to the subunit’s centroid was stored, and the distance along the spline path was calculated to the next protomer.

Twist between protomers along the deformed short-pitch F-actin helix was measured in the context of the moving Frenet-Serret (FS) frame of reference. The FS frame of reference was defined by the orthonormal basis of the unit tangent, unit normal, and unit binormal vectors along the length of the spline. The set of unit tangent vectors sampled at the positions along the 3D cubic spline corresponding to each subunit (as determined during the rise measurements) was calculated. A vector with a magnitude equal to the filament’s radius oriented along the normal axis in the FS frame and its tail at the origin in the FS frame was then rotated in the normal-binormal plane until the distance between its head and the corresponding subunit centroid was minimized. This rotation angle defined the absolute angular twist for the protomer. To measure twist along the short-pitch helix, the difference between consecutive absolute angular twists was calculated.

Inspection of the curvature-twist plots revealed that for bent filaments, there was a clear sinusoidal pattern of the twist measurement if the measurements were separated by protofilament. Furthermore, as the curvature increased along the cryoDRGN trajectory, both the magnitude and position of this sinusoidal pattern changed. Therefore, we modelled the bend-twist phenomenon for each protofilament as travelling waves using the equation:

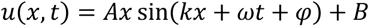

Where u(x,t) is the instantaneous twist, A is a coupling factor between curvature and twist amplitude, k is the propagation factor that determines how rapidly the twist wave travels along the filament’s length with bending, x is the curvature, µ is the period of twist, t is the position along the central axis (parameterized to protomer index), ϕ is the phase shift of twist, and B is the overall average twist. This equation was jointly fit against the measured twist values and estimated curvatures for each of the 16-protomer cryoDRGN models. Curvature for each model was measured as the average of the instantaneous 3D curvatures of the central axis spline. For the ADP nucleotide state, all frames were used. For the ADP-P_i_ state, the first three frames had low curvature and had fluctuating curvature measurements, so they were omitted due to the inaccurate average curvature measurement of the central spline. The fit values for the model parameters are presented in Supplementary Table 3, and example fit functions through experimental data are presented in Supplemental Fig. S5B.

### Analysis of central axis deformations

Analysis of the filaments’ bending deformations were performed on the 16-protomer cryoDRGN models’ central axes. Principal component analysis was performed on the coordinates sampled along the axis spline in Euclidean space. The plane defined by the first and second principal components represents the plane of maximum filament curvature. The plane formed by the first and third principal components represents an orthogonal plane capturing the three-dimensional character of the bent filament. Central axis deformation was also analyzed by measuring the deviation of the central axis from a straight line fit. For each curved cryoDRGN model, a straight line was aligned to the terminal 56 Å of its central axis at the barbed end. Then, discrete sampling steps of size 0.28 Å were made along the central axis, and the distance from the sampled point and the straight line was plotted in Supplementary Fig. 4d.

### Actin subdomain measurements

Actin subdomains were defined using previously established residue assignment conventions^34^: subdomain 1 (SD1): AA 5-32, 70-144, 338-375; SD2: AA 33-69; SD3: AA 145-180, 270-337; SD4: AA 181-269. Using a custom python script, the Euclidean distances, angles, and dihedral angles between subdomains indicated in Supplemental Fig. 5a were measured. The protomer indexing started at the pointed end and progressed to the barbed end. For measurements that spanned multiple protomers, the protomer index corresponded to the most pointed-end protomer.

### Subunit shearing measurements and shear index

For shear measurements, each protomer of the asymmetric F-actin models was aligned to the protomer of the corresponding high-resolution, helically-symmetric model of the same nucleotide state. The average displacement vector for each subdomain between these models was then computed. Observing anti-correlated displacements between non-adjacent subdomains led us to define two shear indices to describe these coordinated deformations: shear index 1, the dot product of the subdomain 1 and 4 displacement vectors, and shear index 2 the dot product of the subdomain 2 and 3 displacement vectors. Shear indices of pairs of subdomain displacement vectors that have large individual magnitudes and opposing directionality will have large negative values, indicative of shear, while small displacements or lack of correlated subdomain displacements will produce values near zero.

### Strain analysis

To quantify protein deformations not explained by rigid-body motions, strain analysis was performed with a custom python script, implementing a previously described approach^35,36^. Briefly, a reference helical F-actin protomer was rigid-body fit into each protomer of the model to which it was being compared. Then, the local deformation matrix within an 8 Å neighborhood was estimated for each alpha-carbon of the reference protomer. The Eulerian strain tensor is computed using a first order approximation of the deformation matrix’s spatial derivative. The shear strain energy is then calculated directly from this strain tensor. This approach to protein deformation has the major advantage of being rotationally invariant and distinguishing rigid-body motions from internal deformation. However, the local deformation estimation can be inaccurate for very large deformations, which limited our strain analysis to individual protomers. Furthermore, the first-order approximation assumes continuous, as opposed to granular, deformations, which makes the measurements relative pseudo-energies.

### Plotting, statistical analysis, and molecular graphics

Plots were generated with GraphPad Prism or Matplotlib^37^. Statistical tests were performed with GraphPad Prism. Molecular graphics were prepared with UCSF Chimera^23^ and UCSF ChimeraX^29^.

